# Anti-TRAP/SSP2 monoclonal antibodies can inhibit sporozoite infection and enhance protection of anti-CSP monoclonal antibodies

**DOI:** 10.1101/2021.10.15.464611

**Authors:** Brandon K. Wilder, Vladimir Vigdorovich, Sara Carbonetti, Nana Minkah, Nina Hertoghs, Andrew Raappana, Hayley Cardamone, Brian G. Oliver, Olesya Trakhimets, Sudhir Kumar, Nicholas Dambrauskas, Silvia A. Arredondo, Nelly Camargo, Stefan H.I. Kappe, D. Noah Sather

## Abstract

Vaccine-induced sterilizing protection from infection by *Plasmodium* parasites, the pathogens that cause malaria, will be essential in the fight against malaria as it would prevent both malaria-related disease and transmission. Stopping the relatively small number of parasites injected by the mosquito before they can migrate from the skin to the liver is an attractive means to this goal. Antibody-eliciting vaccines have been used to pursue this objective by targeting the major parasite surface protein present during this stage, the circumsporozoite protein (CSP). While CSP-based vaccines have recently had encouraging success in disease reduction, this was only achieved with extremely high antibody titers and appeared less effective for a complete block of infection (i.e. sterile protection). While such disease reduction is important, these and other results indicate that strategies focusing on CSP alone may not achieve the high levels of sterile protection needed for malaria eradication. Here, we show that monoclonal antibodies (mAbs) recognizing another sporozoite protein, TRAP/SSP2, exhibit a range of inhibitory activity and that these mAbs can augment CSP-based protection despite conferring no sterile protection on their own. Therefore, pursuing a multivalent subunit vaccine immunization is a promising strategy for improving infection-blocking malaria vaccines.

## Introduction

The last few years have marked a disheartening milestone as the first period in a generation without a reduction in the global burden of malaria ^1^. The interventions that have provided much of the previous progress, such as insecticide-treated bednets and large-scale treatment programs, are highly susceptible to interruptions due to political or economic instability. This was starkly illustrated by the resurgence of malaria in Venezuela in recent years after near-elimination; and in 2020, more globally, due to interruptions in eradication efforts during the COVID-19 pandemic ^1^. Therefore, it is likely that durable, infection-blocking interventions (e.g. vaccines, long-lasting mAbs or chemoprophylactics) will be required to drive malaria to elimination.

Developing such an intervention is hampered by the complex life cycle of the parasite, which begins when an infected mosquito injects tens to hundreds of the “sporozoite” forms of the parasite into the dermis ^2^. From here, sporozoites actively traverse through multiple cell types in search of an endothelial cell through which they will gain access to the blood ^3^. Upon entering the bloodstream, a sporozoite is carried to the liver within minutes, where it traverses multiple cell types in the liver parenchyma and eventually establishes infection in a hepatocyte ^4^. Following ∼7–10 days of development and genome replication (∼2 days in rodent malaria models), each successful liver stage releases 30,000–50,000 “merozoites” that cyclically infect, replicate within and lyse red blood cells ^5, 6^. It is only during this blood stage of infection when symptomatic disease occurs, and is also where a subset of sexually differentiated parasite forms can be picked up by a new mosquito host to continue the transmission cycle. Each step in the infection cycle presents opportunities for intervention, although vaccines targeting the “pre-erythrocytic” stages in the skin and liver have yielded the most promising results ^7^ .

The most advanced pre-erythrocytic vaccine is RTS,S ^8^—an antibody-eliciting subunit vaccine targeting the major sporozoite surface protein circumsporozoite protein (CSP), which has been recently recommended by the WHO ^1, 9^. Vaccines based on attenuated live sporozoites that arrest in the liver and function by a combination of T cells and antibodies have also demonstrated robust protection ^10^. Unfortunately, despite significant efficacy from both approaches in controlled human malaria infection (CHMI) studies in malaria-naive volunteers, both vaccines have markedly reduced efficacy in field trials and have not met the goals of 75% protection against clinical disease for one year as expressed by the WHO ^11^. Recent encouraging Phase II results with the R21 CSP particle in Matrix-M adjuvant do meet this goal ^12^. However, protection with R21 appears dependent on high antibody titers, which would require yearly boosters that are vulnerable to interruptions, and protection is less robust against *infection*. If a vaccine or other intervention (e.g. a mAb or an injectable chemoprophylactic) is to be used as a tool to achieve malaria eradication, it will likely need at least 80% efficacy against *infection* to have a significant and sustained impact on transmission ^13–15^. These realities highlight the significant room for improvement in both T cell and antibody-eliciting vaccines, with the latter more amenable to iterative improvement due to available *in vitro* and *in vivo* preclinical assays ^16–19^.

Of the hundreds of proteins expressed at the sporozoite stage, at least 47 are surface-exposed ^20–22^ and therefore potentially accessible to antibodies. However, few of these proteins have been rigorously investigated for their use in antibody-eliciting vaccines ^23^. In addition to CSP, the thrombospondin-related anonymous protein (TRAP, also known as sporozoite surface protein 2 or SSP2) has been pursued as a vaccine candidate. Similar to CSP, TRAP is essential for sporozoite infectivity ^24, 25^, antibodies against it correlate with protection ^26, 27^ and the protein is abundant ^21^ during the skin stage when parasites are particularly susceptible to antibody mediated inhibition. The TRAP ectodomain consists of 3 main domains: a von Willebrand factor A-like domain (vWA), the thrombospondin repeat (TSR) domain and a repeat domain ^28^. The most advanced TRAP vaccine candidate is an adenovirus/MVA-vectored vaccine eliciting strong T cell responses that has had low or mixed efficacy results in CHMI trials ^29, 30^ and field trials ^31^ but has been improved in mice following targeting of the T cell response to the liver ^32^. Antibody function in experiments involving immunization with TRAP-derived peptides have yielded mixed results ranging from significant sporozoite inhibition in vitro ^33^ to no protection in vivo ^34^ . A combination protein TRAP/RTS,S immunization failed to show significant protection in a clinical trial ^35^, while a fusion-protein approach using TRAP and CSP resulted in complete protection in a small mouse study ^36^. These results using TRAP alone or in combination with CSP are difficult to interpret due to the diversity of vaccine platforms used, the possibility of immune interference in studies combining platforms, and the unclear dominance of roles for antibodies and T cells in protection ^37^. Whether a more targeted TRAP antibody response could contribute to protection either alone or in combination with CSP remains poorly defined.

Here, we used both active immunization and passive transfer of mAbs raised against either *Plasmodium yoelii* (rodent malaria) or *Plasmodium falciparum* (human malaria) TRAP to more directly explore the potential efficacy of anti-TRAP antibodies. We found that anti-TRAP antibodies modestly prevent liver infection in a manner dependent on the TRAP domain recognized. Importantly, we also demonstrate a proof-of-concept that anti-TRAP antibodies with minimal protective capacity of their own can augment anti- CSP antibodies, providing additive protection that raises their protective efficacy above 80% sterile protection. Together, these findings argue for further investigation of rationally designed multi-antigen, antibody-eliciting malaria vaccines or mAb prophylactics that target multiple antigens and might include CSP as well as non-CSP targets such as TRAP.

## Results

### PyTRAP polyclonal antibodies can prevent parasite infection of hepatocytes in vitro and in vivo

To elicit potentially functional anti-TRAP antibodies, we generated full-length ectodomains and fragments of both rodent (*P. yoelii*) and human (*P. falciparum*) malaria TRAP proteins (**Fig. 1A; Suppl. Table 1**) and verified their purity (**Fig. 1B**). Serum from mice immunized with the rodent malaria *P. yoelii* TRAP ectodomain (PyTRAP) recognized *Py* sporozoites by immunofluorescence in a pattern consistent with micronemal localization, indicating the antigenic fidelity of the recombinant protein (**Fig. 2A**). We further tested this serum in an inhibition of sporozoite cell traversal and invasion (ISTI) assay. Compared to control serum, anti-PyTRAP serum was able to modestly but significantly (p=0.028) reduce sporozoite invasion of Hepa1-6 hepatoma cells *in vitro* at a level similar to serum from mice immunized with the recombinant PyCSP ectodomain, although the latter failed to reach significance (p=0.106) (**Fig. 2B**). In contrast, sporozoite traversal of Hepa1-6 cells was not affected by anti-PyTRAP serum (p=0.125), whereas anti-PyCSP serum did significantly reduce traversal (p=0.0057) (**Fig. 2C**). The known inhibitory anti-PyCSP mAb 2F6 ^38, 39^ reduced both inhibition and traversal in this assay, as expected (**Fig. 2B and C**).

**Figure 1.**
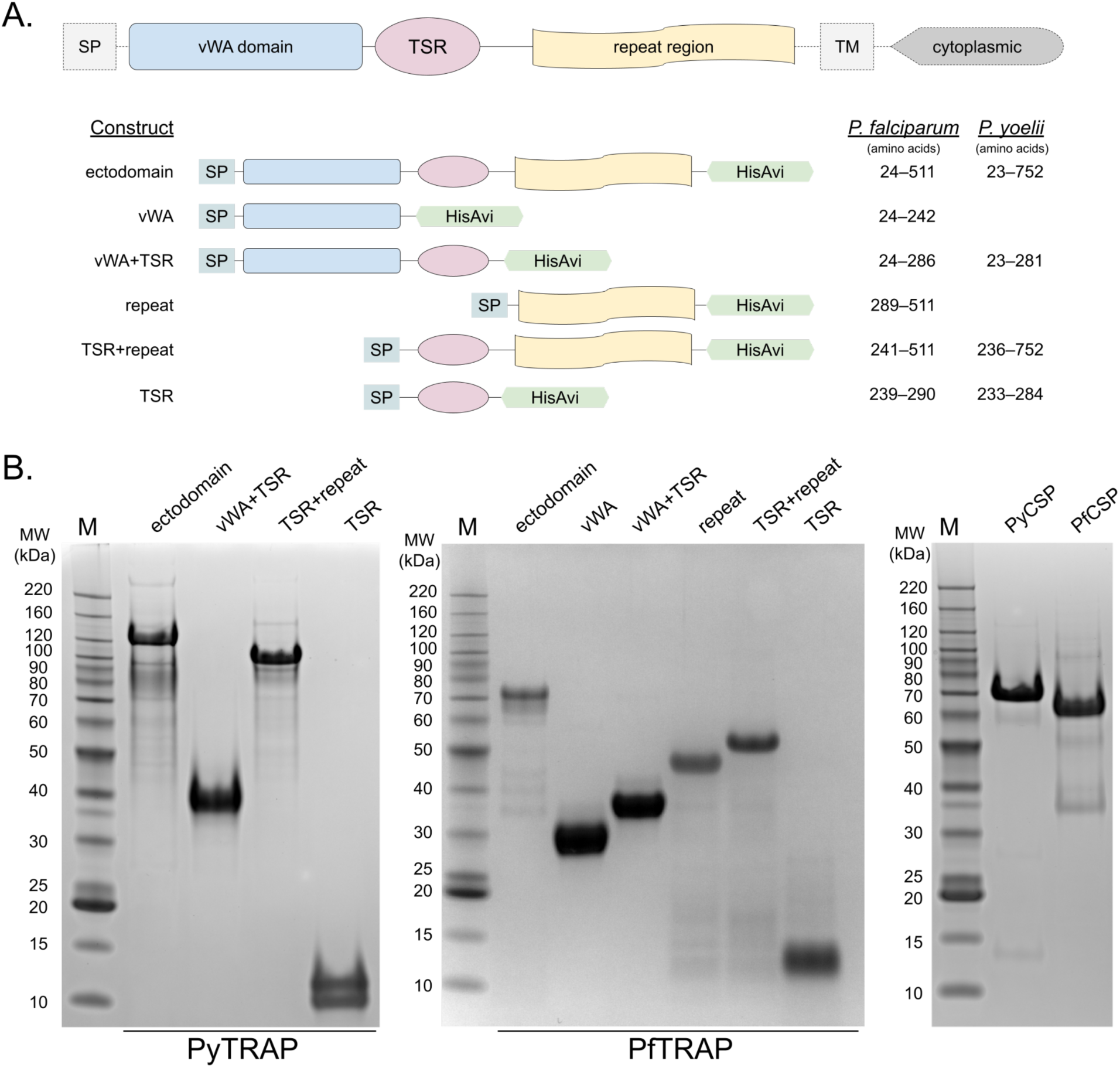
TRAP domain organization and constructs used. Ectodomain and deletion constructs for PyTRAP and PfTRAP generated using the domain boundaries (**A**) were recombinantly expressed and purified alongside the control CSP ectodomain proteins (**B**).

**Figure 2.**
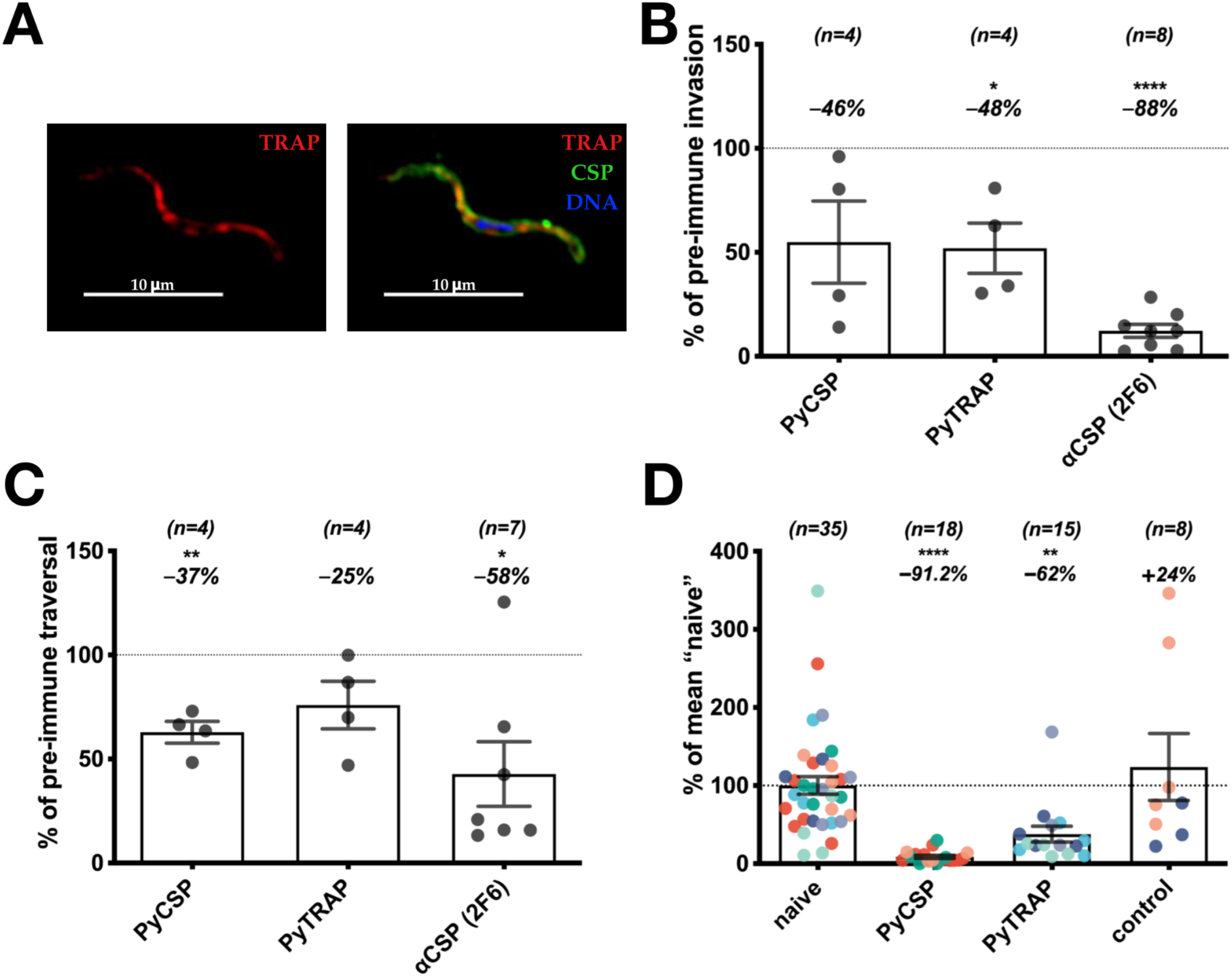
Polyclonal antibodies to PyTRAP inhibit parasite invasion, traversal and in vivo infection. Mice were immunized three times with PyTRAP or PyCSP ectodomains. (**A**) Immune sera were used to verify binding to *Py* sporozoites via immunofluorescence. Shown is a representative example of fixed, permeabilized sporozoites labeled with a 1:100 dilution of polyclonal mouse serum from PyTRAP immunization. The anti-mouse IgG (anti-TRAP serum) is in the red channel shown alone on the left and in combination with anti-CSP mAb 2F6 (green channel) and a DAPI nuclear stain (blue channel); 10-µm scale bars are shown. Immune sera were then assessed for function in vitro for inhibition of invasion (**B**) and traversal (**C**). In **B** and **C**, pooled serum from cohorts of n=5 mice (number of cohorts indicated above each bar) was tested in three independent assays. Each data point represents the average “% of pre-immune” invasion or traversal of these independent assays for each cohort pool. Each bar indicates the group mean, with error bars representing standard error of the mean. Values representing percent changes from 100% (indicated by dotted lines) are shown above. Asterisks indicate a significant difference from 100% as determined by a two-tailed one-sample *t*-test. (**D**) Immunized mice were challenged by the bite of 15 PyGFPluc-infected mosquitos and assessed for parasite liver burden by bioluminescent imaging. Each data point represents an individual mouse with each color corresponding to an independent immunization-challenge experiment (total number of animals shown above each bar). Each data point was normalized to the mean flux from “naive” mice within each challenge experiment, while “control” mice were an additional group immunized with HIV Env gp120 protein. Each bar indicates the group mean, with error bars representing standard error of the mean. Values representing percent changes from 100% (indicated by a dotted line) are shown above. Asterisks indicate significance as determined by ANOVA with Kruskall-Wallis post-test. For **B-D**, * is p≤0.01; and **** is p≤0.0001

PyTRAP-immunized mice were then challenged with *Py* sporozoites via mosquito bite to determine if these antibodies could function *in vivo* to reduce liver infection. We utilized a PyGFPluc parasite, which expresses luciferase, enabling the measurement of liver stage parasite burden by bioluminescence imaging. Mice immunized with a non-specific control protein (Env) showed no reduction in parasite liver stage burden following challenge compared with naive mice. In contrast, mice immunized with the PyTRAP ectodomain showed a significant 62% reduction of parasite liver stage burden. Mice immunized with PyCSP ectodomain showed a 91% reduction relative to naive controls (**Fig. 2D**). Together, these data indicate that anti-PyTRAP antibodies can function *in vitro* and *in vivo* to reduce parasite infection of hepatocytes.

### PyTRAP mAbs display a diverse array of functions in vitro and can provide additive protection to anti-CSP antibodies in vivo

Serum polyclonal antibodies, as studied above, are a mixture of many antibody specificities, making it difficult to characterize the relative contribution to functional activity of responses directed at different domains. To enable such a characterization of the repertoire of PyTRAP-elicited antibodies, we produced a panel of 15 mAbs. When tested in ISTI at 10, 50 and 100 μg/mL, 12 of these mAbs significantly inhibited invasion or cell traversal at one or more concentrations, with mAbs TY03 and TY11 showing the most consistent and potent inhibition (**Fig. 3A**, **Suppl. Fig. 1**).

**Figure 3.**
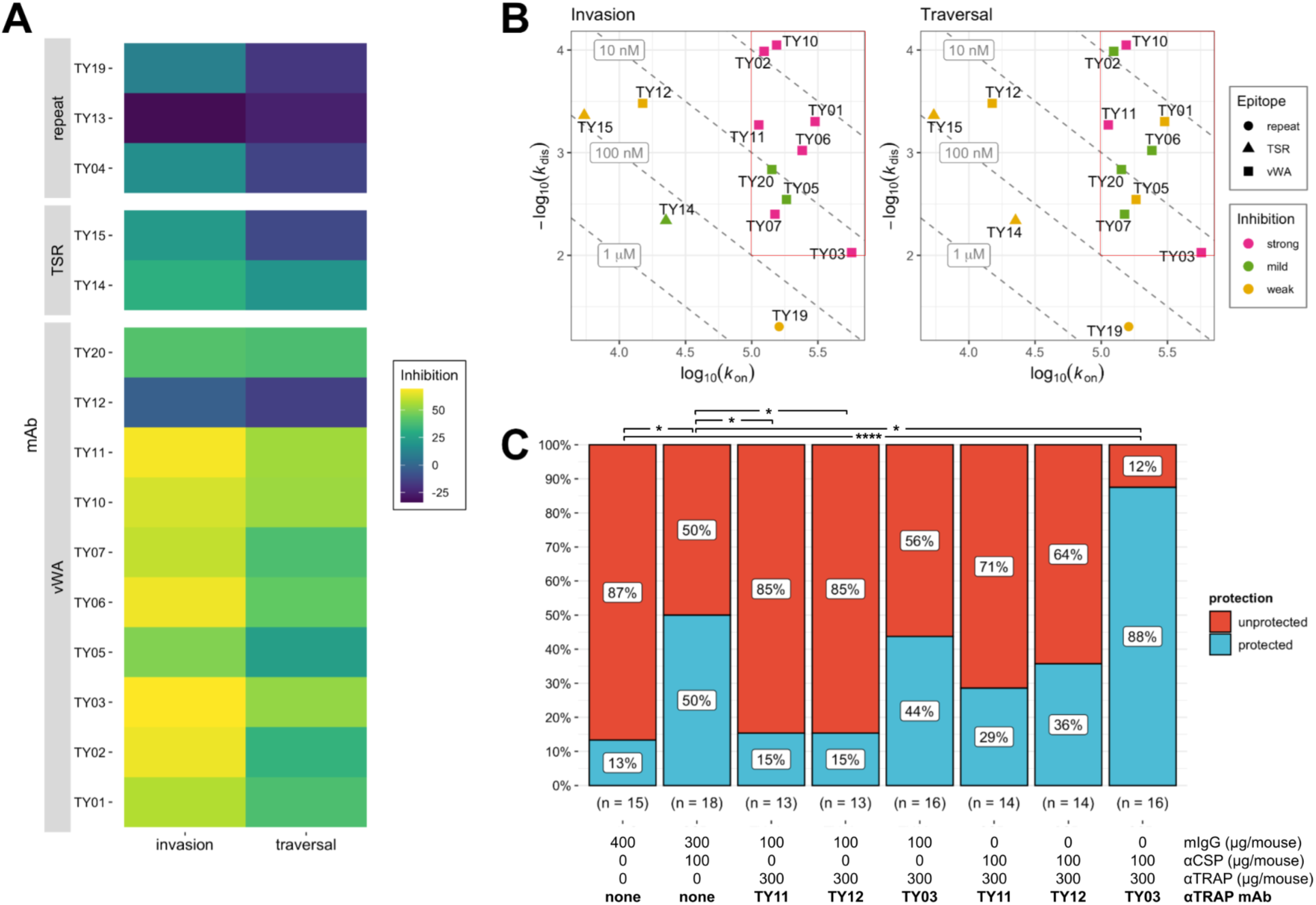
Effects of PyTRAP monoclonal antibodies on parasite activity. (**A**) Each mAb was assessed for in vitro function of inhibition of invasion and traversal. In each case, mean values of % inhibition (i.e., 100% – invasion or traversal value) from the 100-µg/mL mAb concentrations (bar plots with these and additional conditions shown in **Suppl. Fig. 1**) are represented on a color axis. (**B**) Binding kinetics for each mAb was measured by BLI and shown as kinetic maps with gray dashed diagonal contour lines labeled with the corresponding K_d_ values and symbols representing the characterized epitopes for invasion (*left*) and traversal (*right*) inhibition. Higher-affinity (i.e., those possessing lower K_d_ values) mAbs are closer to the upper-right corner of this plot. Symbol color coding represents “strong” inhibition for mean values ≤50%, “mild” inhibition for values ≤70% observed at the 100-μg/mL concentration. Red box highlights the region of the kinetic plots containing the values for mAbs that showed strong inhibition in invasion and traversal assays (**C**) Summarized sterile protection ratios following passive-transfer-challenge experiments (number of animals in each group is shown below the corresponding bar, individual values shown in Table 1). For **C,** * is p≤0.05 and **** is p≤0.0001; values reported were not adjusted for multiple comparison due to small group size and limited comparisons.

**Table 1.**
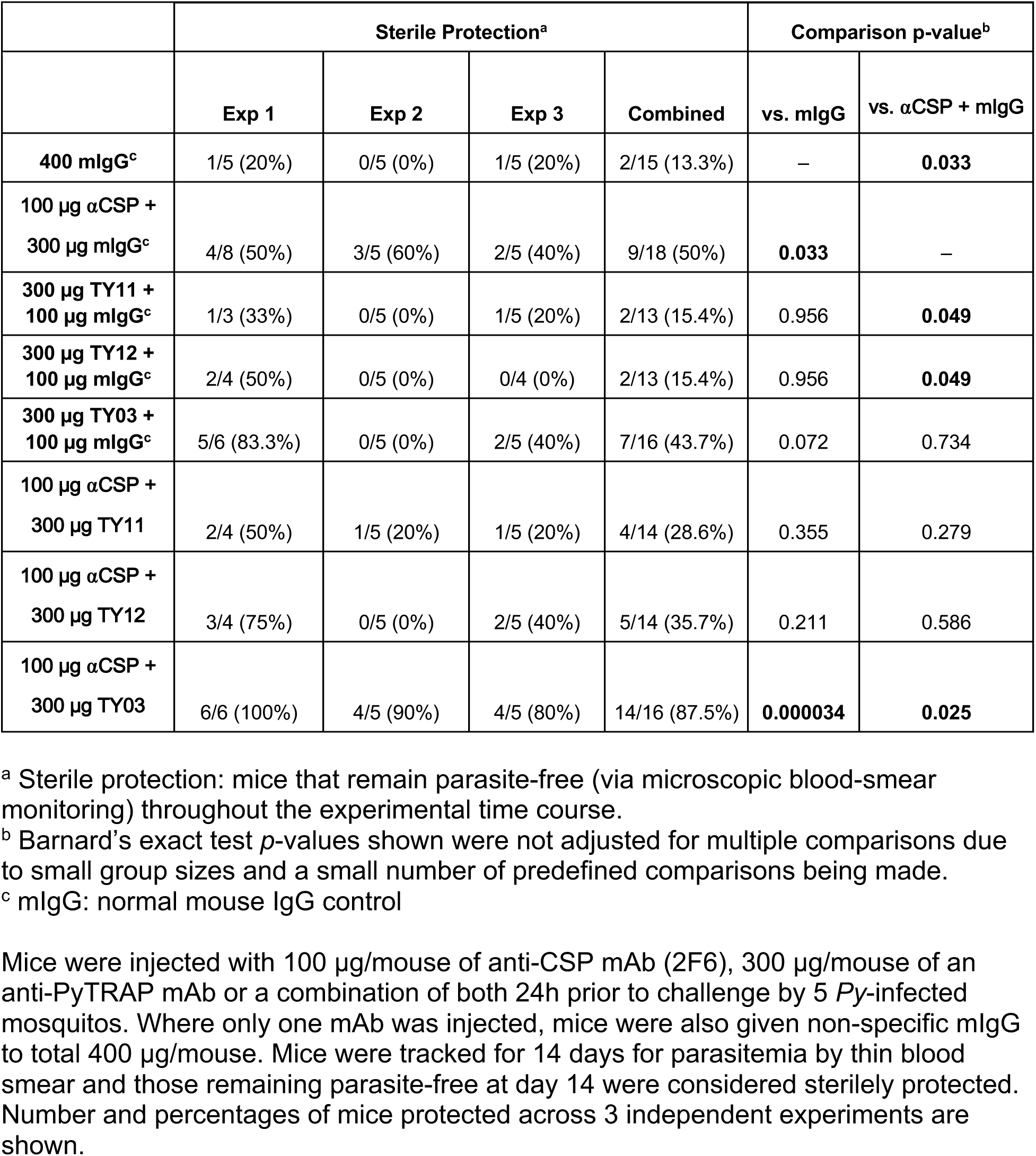
Combination of anti-PyCSP and anti-PyTRAP can improve sterile protection from mosquito bite challenge.

Overall, the mAbs demonstrated a wide range of binding affinities to recombinant PyTRAP (**Fig. 3B**, **Suppl. Table 2**) and recognized epitopes in the vWA, TSR and repeat regions (**Suppl. Table 3**), thus covering the entire protein ectodomain. Among the 15 mAbs recovered, 10 mAbs bound to the vWA domain. Six of these (TY02, TY05, TY06, TY10, TY11, TY20) shared variable-segment assignments for both heavy and light chains, had closely related complementarity-determining-region (CDR) sequences and had 88.4–96.7% and 93.9–96.9% sequence identity in the variable-region sequences of their heavy and light chain, respectively (**Suppl. Table 4 and Suppl. Fig. 2A,B**). As expected, these antibodies were functionally similar in that they bound specifically to the vWA domain (**Suppl. Table 3**), clustered in the same epitope bin (**Suppl. Fig. 3A,B**) and inhibited sporozoite infection *in vitro* (**Fig. 3A, Suppl. Fig. 1**). Two mAbs specifically recognized the TSR domain, and the remaining three mAbs bound epitopes in the repeat region (**Suppl. Table 3**). These non-vWA antibodies showed only modest or no sporozoite inhibition of infection *in vitro* (**Fig. 3A**). Binding interference experiments allowed for the assignment of several distinct epitope bins (**Suppl. Fig. 3A,B**) in addition to the one largely formed by the aforementioned group of mAbs sharing high sequence identity. This likely indicates that the mAbs in our panel bind several distant epitopes on PyTRAP. In addition, this panel of mAbs showed a wide range of binding kinetics, with all strongly inhibitory mAbs having a *k_on_* of >10^5^ M^- 1^s^-1^ and a *k_dis_* of <10^-2^ s^-1^ (**Figure 3B**, note the red box, **Suppl. Table 4**). Together, these data demonstrate that, similar to polyclonal antibodies, anti-PyTRAP mAbs can mediate anti-parasitic function *in vitro*, and that inhibitory function likely depends on fast and stable binding to the vWA domain. However, within the vWA domain some epitopes show higher correlation between binding and blocking of infection compared to others.

We next wanted to determine whether an anti-PyTRAP mAb could provide sterilizing protection *in vivo* on its own or in combination with an anti-CSP mAb. For this, we chose three vWA domain-binding anti-PyTRAP mAbs from distinct epitope bins: TY03 and TY11, which were the top-performing mAbs in ISTI, and TY12, which failed to demonstrate efficacy in ISTI. Neither the anti-PyTRAP mAbs nor the anti-CSP mAb showed significant binding to the mismatched Ag in vitro (**Suppl. Fig. 4A, B**), indicating target specificity. The anti-PyTRAP mAbs were given at 300 μg/mouse (∼15 mg/kg) alone or with a partially protective dose of 100 μg/mouse (∼5 mg/kg) of anti-PyCSP mAb 2F6 prior to mosquito bite challenge ^38^. As shown in **Fig. 3C** and **Table 1**, mice administered anti-PyCSP mAb 2F6 showed significant sterile protection, with 9/18 (50%) remaining blood stage parasitemia-free, compared to 2/15 (13.3%) for mice receiving non-specific murine IgG (p=0.032; this value was not corrected for multiple comparisons due to small sample size and a small number of predefined comparisons being made). Neither TY11 nor TY12 showed any protection (2/13 or 15.4% non-infected) despite TY11 demonstrating the most robust inhibition *in vitro*. Administration of the mAb TY03 resulted in 7/16 mice (43.7%) remaining parasitemia-free, not reaching statistical significance. When combined with the anti-CSP mAb, only the addition of TY03 afforded significant sterile protection (87.5% or 14/16 mice) over the control group (p<0.001), which, importantly, was a significant improvement over protection observed with anti-PyCSP mAb alone (p=0.025; again not corrected for multiple comparisons as above). Together these data indicate that while *in vitro* testing of mAbs can be useful for identifying non-functional mAbs (e.g. TY12), they should be validated *in vivo* for function. Importantly, these data provide proof of concept that non-CSP antibodies can provide additive protection to anti-CSP antibodies.

### Antibodies targeting the human malaria parasite P. falciparum TRAP can function against sporozoite invasion of hepatocytes

We next wanted to determine if antibodies directed against TRAP/SSP2 from the human malaria parasite, *P. falciparum*, could also function to prevent sporozoite infection. Serum from mice immunized with the ectodomain of *P. falciparum* TRAP (PfTRAP) was able to recognize *Pf* sporozoites in IFA (**Fig. 4A**) and demonstrated consistent inhibition of *Pf* sporozoite invasion *in vitro* at a level similar to serum from mice immunized with the ectodomain of *P. falciparum* CSP (PfCSP) (**Fig. 4B**). Inhibition of sporozoite traversal *in vitro* was more modest as compared to anti-PfCSP polyclonal serum (**Fig. 4C**). The known inhibitory anti-PfCSP mAb 2A10 ^40^ demonstrated robust inhibition of both invasion and traversal (**Fig. 4B and C**).

**Figure 4.**
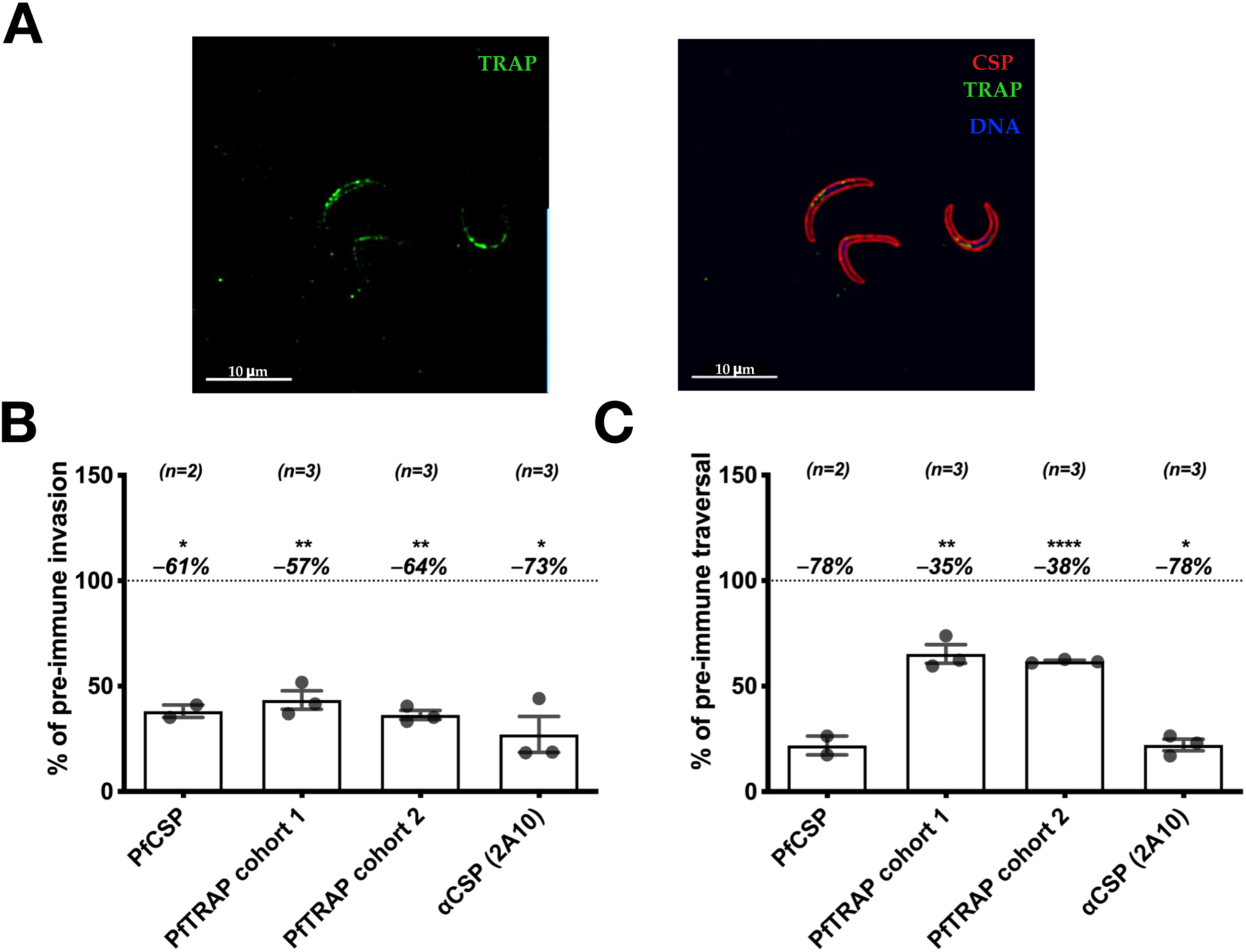
Polyclonal antibodies to PfTRAP inhibit parasite invasion and traversal in vitro. Mice were immunized three times with PfTRAP or PfCSP ectodomains. (**A**) Immune sera were used to verify binding to *Pf* sporozoites via immunofluorescence. Shown are fixed, permeabilized sporozoites labeled with a 1:800 dilution of polyclonal anti-PfTRAP mouse serum (followed by anti-mouse IgG secondary; green channel), fluorescently labeled anti-PfCSP monoclonal antibody 2A10 (red channel, right image) and DAPI nuclear stain (blue channel, right image); 10-µm scale bars are shown. Immune serum was then assessed for function in vitro for inhibition of invasion (**B**) and traversal (**C**). In **B** and **C**, each data point is the average “% of pre-immune” invasion or traversal from technical triplicates in independent experiments; two separate immunization experiment sets are represented as “PfTRAP cohort 1” and “PfTRAP cohort 2”. Each bar indicates the group mean, with error bars representing standard error of the mean and percent change from 100% (shown as dashed line) shown above. Asterisks indicate a significant difference from 100% as determined by two-tailed one-sample *t*-test where * is p≤0.05; ** is p≤0.01; and **** is p≤0.0001

Using a similar approach to the anti-PyTRAP work described above, we isolated 7 anti-PfTRAP mAbs from immunized mice. Of these, 5 mAbs recognized the vWA domain with AKBR-3, AKBR-4 and AKBR-6 likely recognizing adjacent epitopes (**Suppl. Fig. 3C,D**), and 2 mAbs recognized the TSR domains (**Suppl. Table 3**). In contrast to the high proportion of functional anti-PyTRAP mAbs (12 of 15), only 2 of 7 anti-PfTRAP mAbs, both recognizing the vWA domain, showed any sporozoite-inhibitory function *in vitro*: AKBR-4 and AKBR-10. Further, only AKBR-4 demonstrated significant inhibition of both invasion and traversal (**Fig. 5A, Suppl. Fig. 5**), despite having unremarkable binding properties with the PfTRAP ectodomain (**Fig. 5B**). Surprisingly, mAb AKBR-7, which had the best binding properties of the set (K_d_ ∼0.15±0.04 nM, **Suppl. Table 2**), demonstrated the worst inhibitory properties (**Fig. 5B**). Similar to the case with the anti-PyTRAP mAb panel described above, our data suggest that the PfTRAP vWA domain contains epitopes exposing vulnerability to inhibition, however lack of mAbs that strongly bind other portions of PfTRAP make it difficult to discount the roles that these domains may play in inhibition *in vivo*.

**Figure 5.**
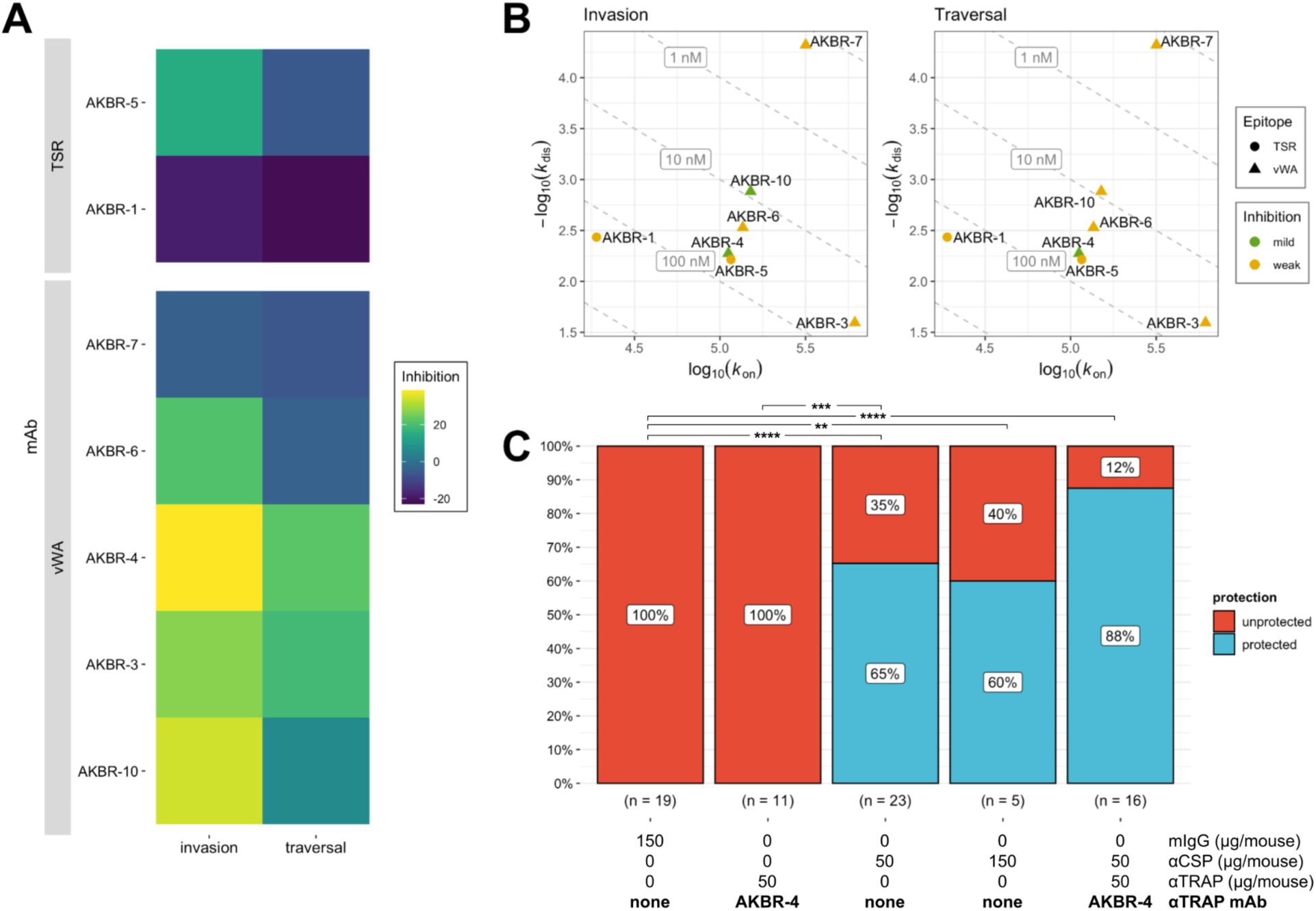
Monoclonal antibodies to PfTRAP inhibit parasite invasion, traversal in vitro. (**A**) Each mAb was assessed for in vitro function of inhibition of invasion and traversal. In each case, mean values of % inhibition (i.e., 100% – invasion or traversal value) from the 100-µg/mL mAb concentrations (bar plots with these and additional conditions shown in Suppl. Fig. 5) are represented on a color axis. (**B**) Binding kinetics for each mAb was measured by BLI and shown as kinetic maps with gray dashed diagonal contour lines labeled with the corresponding K_d_ values and symbols representing the characterized epitopes for invasion (*left*) and traversal (*right*) inhibition. Higher-affinity (i.e., those possessing lower K_d_ values) mAbs are closer to the upper-right corner of this plot. Symbol color coding represents “mild” inhibition for values ≤ 70% and “weak” for mean values >70% observed at the 100-μg/mL concentration. (**C**) Summarized sterile protection breakdowns following passive-transfer-challenge experiments (number of animals in each group is shown below the corresponding bar, individual values shown in Table 2). For **C,** ** is p≤0.01, and **** is p≤0.0001; values reported were not adjusted for multiple comparisons due to small group size and limited comparisons.

**Table 2.** Combination of anti-PfCSP and anti-PfTRAP can improve sterile protection from mosquito bite challenge.

|  | Sterile Protection<sup>a</sup> |  |  |  | Comparison p-value<sup>b</sup> |  |
| --- | --- | --- | --- | --- | --- | --- |
|  | Exp 1 | Exp 2 | Exp 3 | Combined | vs. mlgG | vs. 50 µg αCSP |
| **150 µg mlgG<sup>c</sup>** | 0/7 (0%) # | 0/7 (0%) # | 0/5 (0%) | 0/19 (0%) | – | **<0.0001** |
| **50 µg AKBR-4** | 0/6 (0%) | – | 0/5 (0%) | 0/11 (0%) | 1 | **0.0002** |
| **50 µg αCSP** | 5/7 (71%) # | 5/8 (63%) # | 5/8 (63%) | 15/23 (65%) | **<0.0001** | – |
| **150 µg αCSP** | – | 3/5 (60%) | – | 3/5 (60%) | **0.002** | 0.88 |
| **50 µg AKBR-4 +**<br>**50 µg αCSP** | – | 6/7 (86%) | 8/9 (89%) | 14/16 (88%) | **<0.0001** | 0.131 |

<sup>a</sup> Sterile protection: mice that remain parasite-free (via microscopic blood-smear monitoring) throughout the experimental time course.
<sup>b</sup> Barnard's exact test *p*-values shown were not adjusted for multiple comparisons due to small group sizes and a small number of predefined comparisons being made.
<sup>c</sup> mlgG: normal mouse IgG control

### A vWA-directed anti-PfTRAP mAb increases the protection afforded by a protective CSP mAb

Because *Pf* sporozoites do not infect murine livers, the only means to test the activity of anti-*Pf* antibodies against sporozoite infection *in vivo* is by either challenging passively or actively immunized wild-type mice with transgenic rodent parasites expressing the *Pf* proteins of interest ^41–43^ or by passive immunization of immune-deficient humanized liver mice (FRGhuHep) that can be challenged with wild type *Pf* sporozoites ^17^. We chose to utilize the latter as it is an established model of antibody-mediated protection against *Pf* infection ^17, 44–49^ and allows testing of any future combination of anti-*Pf* antibodies without the need for generating combinatorial transgenic parasites. In this model, humanized-liver mice receive a passive transfer of antibodies and are then infected with *Pf* sporozoites via mosquito bite. Six days later, mice are injected with human red blood cells, which can then be infected by merozoites emerging from the liver, and blood stage infection can be quantified by qRT-PCR on days 7 and 9. In this model, detection of parasites by qRT-PCR on either days 7 or 9 has proven to be a stringent and sensitive means of detecting the presence of blood stage parasites ^17, 50, 51^ . Therefore, we define sterile protection in this model as the absence of parasites in the blood above the limit of detection at either day 7 or 9.

Using this method, we tested the ability of the anti-PfTRAP mAb AKBR-4 to provide sterile protection against *Pf* mosquito-bite infection alone or in combination with a partially-protective anti-PfCSP mAb CIS43 ^45^. Neither the anti-PfTRAP mAb nor the anti-CSP mAb showed significant binding to the mismatched Ag in vitro (**Suppl. Fig. 4C, D**), indicating target specificity. We chose a dose of 50 μg/mouse (∼2.5 mg/kg) for each mAb as this provides partial protection with an anti-PfCSP mAb ^45^ and gives a serum concentration of ∼10 μg/mL at the time of infection, which is achievable by both active vaccination and passive transfer of long-lasting mAbs ^52, 53^. We previously conducted passive administration, mosquito bite challenge in 2 independent experiments ^45^ , which showed that 50 µg/mouse dose of anti-PfCSP mAb CIS43 was protective (5/7 and 5/8 protected in each experiment), compared to control mice (0/7 and 0/7 protected). To avoid unnecessary repetition of FRGhuHep experiments, we included those cohorts in our overall analysis of mAbs in this study and conducted a third independent experiment with the control and 50 µg/mouse dose of anti-PfCSP mAb CIS43 groups in each. In these experiments, 50 µg/mouse mAb CIS43 yielded a total of 15/23 protected (65%), which was significant compared to 0/19 of control mice protected (0%, p<0.0001; **Table 2**, **Fig. 5C**). To determine if the protection afforded by CIS43 would scale linearly with dose and possibly reach 100%, we included a group of 5 FRGhuHep mice in a single experiment, in which the dose was increased 3-fold to 150 μg/mouse. This resulted in 3 of 5 mice protected (60%; p=0.002 over control) but was not significantly different from the groups that received 50 μg/mouse.

On its own, passive administration of 50 μg/mouse of AKBR-4 failed to provide any sterile protection over two of these experiments (0/11, 0%). Yet, when 50 μg/mouse of AKBR-4 was combined with 50 μg/mouse of the anti-PfCSP mAb (100 μg mAb/mouse total), 14/16 (88%; p<0.0001 over control) mice were sterilely protected over two independent experiments. The improvement afforded by the AKBR-4/anti-PfCSP mAb combination over the efficacy of the anti-PfCSP mAb alone trended toward, but did not reach statistical significance at this group size (p=0.131). Together, these results provide the proof of concept that antibodies directed against PfTRAP can reduce *Pf* sporozoite cell traversal and invasion of hepatocytes *in vitro*, and potentially enhance the protection of anti-CSP mAbs when used in combination with the latter.

## Discussion

Studies examining CSP-elicited antibody responses have shown that within a polyclonal antibody population only a subset are highly potent antibody clones, and their distinguishing binding properties can be quite nuanced ^39, 44, 45, 54–58^. Understanding the characteristics associated with protection is crucial for the development of superior mAb products and vaccine immunogens, yet such studies have not been previously performed for TRAP or other non-CSP pre-erythrocytic antibody targets. Here, we show that the polyclonal antibody response to PyTRAP ectodomain can substantially reduce parasite infection of hepatocytes *in vitro* and use mAbs to conclude that this effect is likely driven by vWA and TSR-specific antibodies. These findings are in line with some previous work using antibodies against TRAP protein fragments ^33^, yet they contrast other observations that failed to see significant inhibition ^34^. Our data with PfTRAP were more limited but the only mAb that was functional *in vitro* also recognized the vWA domain. Taken together, our data with polyclonal and monoclonal antibodies clearly demonstrate that TRAP is a viable antibody target, and that its vWA domain contains sites of vulnerability.

Critical for any vaccine or mAb product that can be used for malaria eradication will be achieving high levels of sterile protection at sustainable antibody levels. Experience with RTS,S—which elicits extremely high peak levels of anti-CSP antibodies—as well as published data describing the activity of potent anti-CSP mAbs in animal models suggest that increasing anti-CSP antibody titers can increase protection ^45, 55, 59^. The first CHMI trial using passive transfer of the anti-PfCSP mAb CIS43 (also used in this study) showed that mAbs can provide sterilizing protection against *P. falciparum* mosquito-bite infection at serum concentrations between ∼50–500 µg/mL ^60^ . However, maintenance of such high antibody titers for over a year may not be sustainable for active or passive immunization strategies. As an alternative to frequent vaccine boosting or mAb injections to sustain high titers, it may be possible to achieve high levels of protection at lower antibody titers using multivalent vaccination or multiple mAbs recognizing distinct protein targets. However, there have been no studies to date directly addressing this question, which is best examined using passive transfer of antibodies followed by mosquito bite challenge, as done here.

Our data in the *Py* model show that an anti-PyTRAP mAb, offering no significant protection by itself, can improve protection against mosquito-bite infection of a partially-protective anti-CSP mAb regimen. Our experiments using *Pf* mosquito-bite challenge in FRGhuHep mice, which received a combination of anti-PfCSP and anti-PfTRAP mAbs, did not show a statistically significant improvement over anti-PfCSP mAb treatment alone. However, the fact that this combination was the only regimen to deliver strong protection in repeated experiments, as well as the strong statistical trend observed, offer support for such an approach against *P. falciparum*. Importantly, the 88% sterile protection was achieved using a low total dose of mAb (100 µg/mouse or ∼5 mg/kg). This total dose of 100 µg/mouse (50 µg/mouse each of anti-PfCSP and anti-PfTRAP mAb) is expected to give a total circulating mAb concentration of ∼20 µg/mL ^45^ —a level that can be achieved for ∼36 weeks with a single 20 mg/kg injection of long-lasting mAbs ^52, 60, 61^ or ∼4 years via active vaccination ^62^ . Although it remains to be seen how accurately these animal models translate to the clinic, these data suggest that reaching the 80% sterile protection threshold needed for vaccines ^11^ or injectable anti-malarials ^15^ that can be used as eradication tools may be achieved by targeting multiple proteins rather than by increasing the concentration of antibodies recognizing CSP alone.

Intriguingly, our data showed that some anti-TRAP mAbs, when combined with anti-CSP mAbs, resulted in enhanced protection despite providing no statistically significant sterile protection on their own. These observations may be explained by the fact that the sterile protection readout requires the prevention of all parasites from successfully infecting the liver, effectively introducing a threshold effect. Therefore, it is possible that a weakly inhibitory mAb would have a more pronounced effect in combination with a partially protective regimen (e.g., that of a suboptimal dose of an anti-CSP mAb) than would be predicted by single-mAb experiments when using sterile protection as a readout. Additional studies clarifying the additive vs. synergistic nature of this or any multivalent approach will be needed to determine the utility of combining CSP with other immunogens, but will require large group sizes and experiments designed specifically to test such hypotheses.

In summary, we present a proof-of-concept that antibodies targeting TRAP can contribute to sterile protection when used in combination with anti-CSP antibodies. These findings support vaccine and mAb strategies involving multiple *Plasmodium* pre-erythrocytic-stage antigens, and argue that efforts to develop a long-lasting, infection-blocking malaria intervention would greatly benefit from identifying non-CSP antibody targets that can enhance CSP-elicited protection. Although such a multivalent approach can be achieved with mAbs, it is currently limited by cost ^63^. Active vaccination with multiple antigens has been hampered by challenges of generating and combining multiple protein-in-adjuvant formulations, although this may be more easily achieved by the use of mRNA-based vaccines, which have proven adept as a multi-antigen vaccine platform in preclinical studies ^64, 65^ . Our data, which demonstrate that enhanced protection over CSP-only strategies is possible by way of multivalent subunit vaccination or delivery of mAbs, provide the impetus to pursue such strategies in preclinical studies that better define additive protection and identify additional targets.

## Materials and Methods

### Recombinant protein production

Recombinant proteins were produced in transiently transfected suspension culture of FreeStyle 293 cells (Thermo Fisher Scientific, Waltham, MA, USA), as previously described ^66^. Briefly, codon-optimized constructs encoding the ectodomains or deletion constructs of *Plasmodium falciparum* TRAP (PfTRAP), *Plasmodium yoelii* CSP (PyCSP) and *Plasmodium yoelii* TRAP (PyTRAP) were generated as fusions to the 8xHis and AviTag ^67^ sequences (**Suppl. Table 1**). Following transfection using the high-density PEI method ^68^ and the subsequent 5-day incubation, cells were removed by centrifugation and the culture supernatants were supplemented with NaCl (+350 mM) and sodium azide (0.02%). Treated culture supernatants were passed by gravity through NiNTA agarose, washed with Wash Buffer (10 mM Tris-HCl, pH 8, 300 mM NaCl, 10 mM imidazole), and eluted with Elution Buffer (10 mM Tris-HCl, pH 7.4, 300 mM NaCl, 200 mM imidazole). Further purification was performed by size-exclusion chromatography using a calibrated Superdex 200 (10/600) column (Cytiva, Marlborough, MA, USA). The HIV Env gp120 control protein was produced, as previously described ^69^. When required, site-specific biotinylation with BirA and buffer-exchange by gel filtration was performed, as previously described ^66^.

### Antibody cloning and production

Antibodies were cloned and produced, as previously described ^66^. Briefly, ectodomain PfTRAP and PyTRAP constructs were used as immunogens, and their biotinylated versions were used to isolate antigen-specific B cells by flow cytometry (see sample gating strategy in **Suppl. Fig. 6**). Following culture previously described medium ^66^ modified by the addition of 1.5 µM CpG (ODN-1826) (Integrated DNA Technologies, Coralville, IA, USA), wells containing B cells producing antigen-binding IgG were identified by ELISA, immunoglobulin-encoding transcripts were amplified by RT-PCR and used for the generation of heavy- and light-chain constructs for recombinant mAb expression. The sequences were annotated using IgBLAST ^70^.

To express recombinant mAbs, the plasmid DNA was used to transfect suspension cultures of FreeStyle 293 cells (Thermo), as described above. After five days in culture, cells were removed by centrifugation and the cultures were supplemented with NaCl (+350 mM) and sodium azide (0.02%). Treated culture supernatants were passed by gravity through Protein G resin equilibrated in Wash Buffer (10 mM HEPES, pH 7, 300 mM NaCl, 2 mM EDTA), washed with Wash Buffer, and eluted with 100 mM glycine, pH 2.7. Resulting eluates were buffer-exchanged by repeated centrifugal ultrafiltration with HBS-E (10 mM HEPES, pH 7, 150 mM NaCl, 2 mM EDTA).

### Binding properties of mAbs

Binding kinetics measurements were characterized using biolayer interferometry (BLI) measurements on an Octet QK^e^ instrument (Sartorius, Göttingen, Germany), as previously described ^66^. Briefly, antibodies in culture supernatants were immobilized on anti-Mouse IgG Fc Capture biosensors and allowed to associate with antigen serially diluted (in the range of 1–1000 nM) in 10x Kinetics Buffer (10xKB: PBS + 0.1% Bovine Serum Albumin, 0.02% Tween-20 and 0.05% sodium azide) followed by dissociation in 10x KB. Resulting sensorgram data was evaluated using ForteBio Data Analysis software (version 7.0.1.5) to generate a fit to the 1:1 binding model and provide estimates for the *k_on_* and *k_dis_* rate constants (see sample sensorgrams and fitted curves in **Suppl. Fig. 7**).

The relative specificity of Ag recognition by the mAbs was assayed using biotinylated Ags (30 µg/mL, with the exception of PfCSP, which was used at 10 µg/mL) immobilized on streptavidin biosensors, and incubated with mAbs (50 µg/mL, with the exception of anti-PfCSP, which was used at 10 µg/mL) diluted in 10xKB.

Epitope bins within anti-TRAP mAb panels were assigned based on the interference patterns similar to previous work ^71, 72^. First, His-tagged PyTRAP or PfTRAP (30 µg/mL) was immobilized on NiNTA biosensors in HBS-NPM buffer (20 mM HEPES, pH 7, 150 mM NaCl, 1 mM MgCl_2_, 0.1 mg/mL Bovine Serum Albumin, 0.05% NaN_3_, 0.02% Tween-20). Interference for each pair of mAbs was assessed by binding the first mAb (mAb1) (50 µg/mL, except for TY14 and TY15, which were used at 100 µg/mL) to saturation before allowing the binding from the second mAb (mAb2) (50 µg/mL) to take place. Magnitude of the signal for each mAb2 binding event was corrected by subtracting the signal for the corresponding mAb1 binding step. Additionally, the signal for each mAb2 binding was collected in absence of pre-bound mAb1 (i.e., “blank” HBS-NPM buffer was used in place of mAb1 solution) and used to normalize the corrected mAb2 signal. Finally, the normalized mAb2 values were collected for each mAb1 and the resulting interference pattern sets were used to calculate the Pearson correlation coefficients using R (version 4.0.2) and plotted using the R packages pheatmap (1.0.12). Network graphs were plotted using R package igraph (version 1.2.10) with edges connecting pairs of nodes with a Pearson correlation coefficient >0.7; and clusters of interconnected nodes are referred to as epitope bins.

### Coarse epitope mapping by ELISA

Domain specificity of the mAbs was characterized by enzyme-linked immunosorbent assay (ELISA) using TRAP ectodomain and fragments from PfTRAP and PyTRAP, as previously described ^66^.

### Sporozoite production

For rodent parasite (*P. yoelii*), female Swiss Webster mice for parasite maintenance were purchased from Envigo (Livermore, CA, USA) and injected intraperitoneally (i.p.) with blood stage PyGFPluc ^73^. Three days later, gametocyte exflagellation was confirmed and the infected mice were used to feed female *Anopheles stephensi* mosquitoes. Fourteen to 16 days after the feed, salivary gland sporozoites were isolated from the mosquitoes and used in mouse infections.

For human malaria (*P. falciparum*) experiments, infected *A. stephensi* mosquitoes were produced, as previously described ^74^.

### Animal studies ethics statement

All procedures involving animals were performed in adherence to protocols of the Institutional Animal Care and Use Committee (IACUC) at the Seattle Children’s Research Institute.

### Mouse active immunization and challenge

To generate polyclonal serum and a source of mouse mAbs, six to eight week-old BALBc/J mice were purchased from Jackson Laboratories (Bar Harbor, ME, USA) and injected intramuscularly three times at days 0, 14 and 38 using Adjuplex mixed with 20– 25 µg of target protein. Mice immunized with recombinant *Py* proteins were then challenged by the bite of 15 PyGFPluc-infected mosquitos, as described ^38^. Briefly, the proportion of mosquitos infected with *Py* was determined by presence of midgut oocysts on days 7–12. This proportion was used to prepare a cage with 15 infected mosquitoes per animal (i.e. if 50% of mosquitoes had oocysts, 30 mosquitos/animal were used). These mosquitos were then exposed to anesthetized mice for 10 minutes with lifting of mice every minute to encourage active probing as opposed to blood feeding. Forty-two hours later, parasite liver burden was assessed by bioluminescent imaging, as previously described ^38^. Mice were then immediately sacrificed and splenocytes collected and cryopreserved for B cell isolation and mAb production.

Mice immunized with *Pf* proteins were immunized as above with the exception that mice were additionally boosted with IV protein three days prior to sacrifice and collection and cryopreservation of splenocytes.

For both, serum was collected from immunized mice by collecting whole blood in BD microtainer serum tubes (Becton-Dickinson, Franklin Lakes, NJ, USA), allowing blood to clot at room temperature for at least 30 minutes and then centrifuged according to manufacturer’s instructions to separate serum for storage and use in *in vitro* assays.

### Sporozoite immunofluorescence microscopy

*Py* or *Pf* sporozoites were stained as previously described ^75^. Briefly, freshly dissected *Py* or *Pf* sporozoites were fixed with 4% PFA and air-dried onto glass slides overnight. These were then permeabilized with 0.1% Triton X-100 and stained with polyclonal (serum at 1:100-1:800 dilution) or monoclonal (10 µg/mL) antibodies. Sporozoites were identified by co-staining with anti-CSP mAbs as well as DAPI for nuclear localization. Images were acquired using an Olympus IX-70 DeltaVision deconvolution microscope at 100X magnification.

### In vitro inhibition of sporozoite traversal and invasion (ISTI)

*In vitro* ISTI was performed, as previously described for both *Py* and *Pf* ^76^. Briefly, freshly-dissected sporozoites were added to hepatoma cells (Hepa1-6 for *Py* and HC-04 for *Pf*) plated a day prior in 96-well plates in the presence of antibodies and FITC-dextran in technical duplicates or triplicates. After 90 minutes, cells were fixed, permeabilized and stained with Alexa Fluor 647-labeled anti-CSP mAbs and analyzed by flow cytometry. Invaded cells were identified by the presence of CSP and traversed cells by the uptake of FITC-dextran with gating set to uninfected, stained wells. Within each experimental replicate, antibody-treated wells were normalized to the invasion and traversal of wells treated with pre-immune serum or non-specific mouse IgG, which was set to 100%.

### Anti-Py mAb passive transfer and challenge

Six to eight week-old BALBc/J mice were intravenously injected with indicated doses of mAbs 24 hours prior to challenge by bite of 5 PyGFPluc-infected mosquitos following the same methods as described above for “Mouse active immunization and challenge”. Mice were followed up for infection by Giemsa-stained thin blood smear every other day from days 3–14 for identification of blood stage parasites. Mice in which we failed to identify parasites in 40,000 red blood cells over the entire period were considered negative and sterilely protected. Control mice were administered non-specific polyclonal mouse IgG at a dose equivalent to the highest dose in experimental groups.

### anti-Pf mAb passive transfer in FRG humanized liver mice

Mice repopulated with human hepatocytes (FRGhuHep) were purchased from Yecuris, Inc. (Tualatin, OR, USA) and infected with *Pf* via mosquito bite, as previously described ^17, 45^. Briefly, indicated doses of mAb were intravenously injected into mice 24 hours prior to challenge by the bite of 5 *Pf*-infected mosquitos using the same criteria and methods as above. On day 6 post-infection, mice were intravenously injected with 400 µL of human red blood cells. On days 7 and 9 post-infection, 100 µL of peripheral blood was collected, immediately added to Nuclisens Buffer (bioMerieux, Inc., Durham, NC, USA) for lysis and storage. Parasite presence was quantified by qRT-PCR for *Pf* 18s rRNA, as previously described ^77^. Any mouse with a Ct value above the “no template” control at either day was considered positive for parasitemia.

### Statistics

Statistical analyses and plotting were carried out in Prism (version 9.2.0) (GraphPad Software, San Diego, CA, USA) or in R (version 4.0.2) using packages Exact (version 2.1), ggpubr (version 0.4.0), ggstatsplot (version 0.7.2). Statistical tests and outcomes are noted in the figure legend for each figure. For all tests, a p-value of <0.05 was considered significant and values not specifically labeled were above this threshold.

## Data availability

DNA sequences encoding the mAbs described here have been deposited in GenBank (accession numbers OK484322–OK484365).

## Acknowledgements

We would like to thank the vivarium staff at Seattle Children’s Research Institute for their support of animal studies, and Weldon DeBusk for his assistance with the flow cytometry experiments. We would like to thank Dr. Sean C. Murphy of the University of Washington for performing the *Pf* qPCR analysis. Additionally, we would like to thank Dr. Paul T. Edlefsen of the Fred Hutch Cancer Research Center for the helpful discussions of statistical analysis and Drs. Nevile Kisalu and Robert Seder of the NIH VRC for their provision of mAb CIS43. This study was funded by NIH R01 AI117234 to S.H.K. and D.N.S.

## Author contributions

B.K.W. and V.V. contributed equally to this work.

Conceptualization and experimental design: B.K.W., V.V., S.H.I.K. and D.N.S.

Investigation: B.K.W., V.V., S.C., N.M., N.H., A.R., H.C, B.G.O., O.T., S.K., N.D., S.A.A. and N.C.

Data analysis and visualization: B.K.W., V.V., N.H., N.M.

Writing — Original draft: B.K.W.

Writing — Review and editing: B.K.W., V.V., S.H.I.K. and D.N.S.

Resources: S.H.I.K. and D.N.S.

Supervision, Project Administration and Funding Acquisition: B.K.W., S.H.I.K., D.N.S.

