## Supplemental Tables and Figures for "Anti-TRAP/SSP2 monoclonal antibodies can inhibit sporozoite infection and enhance protection of anti-CSP monoclonal antibodies"

#### **Supplementary Information**

*Supplementary Table 1. Constructs used.*

| Protein (Database Reference) | Domain(s) | Amino Acids |
|---|---|---|
| PyTRAP (PlasmoDB:PYYM_1351500) | Ectodomain | 23–752 |
| | vWA+TSR | 23–281 |
| | TSR | 233–284 |
| | TSR+Repeats | 236–752 |
| PfTRAP (PlasmoDB:PF3D7_1335900) | Ectodomain | 24–511 |
| | vWA+TSR | 24–286 |
| | vWA | 24–242 |
| | TSR | 239–290 |
| | TSR + Repeats | 241–511 |
| | Repeats | 289–511 |
| PyCSP (PlasmoDB:PYYM_0405600) | Ectodomain | 21–362 |
| PfCSP (PlasmoDB:PF3D7_0304600) | Ectodomain | 21–374 |

*Supplementary Table 2: Binding kinetics for anti-TRAP mAbs.*

| mAb name | epitope[&] | target | $K_D$ (M)[*] | $k_{on}$($M^{-1}s^{-1}$)[*] | $k_{on}$ Error[*] | $k_{dis}$($s^{-1}$)[*] | $k_{dis}$ Error[*] | Full $\chi^2$[*] | Full $R^2$[*] |
|---|---|---|---|---|---|---|---|---|---|
| AKBR-1 | TSR | PfTRAP | 1.94E-07 | 1.90E+04 | 1.42E+03 | 3.68E-03 | 1.84E-04 | 0.038749 | 0.950903 |
| AKBR-3 | vWA | PfTRAP | 4.15E-08 | 6.14E+05 | 1.36E+05 | 2.55E-02 | 1.25E-03 | 0.213402 | 0.947356 |
| AKBR-4 | vWA | PfTRAP | 4.76E-08 | 1.12E+05 | 4.60E+03 | 5.31E-03 | 7.55E-05 | 0.021718 | 0.986626 |
| AKBR-5 | TSR | PfTRAP | 5.29E-08 | 1.16E+05 | 4.52E+03 | 6.11E-03 | 6.67E-05 | 0.069659 | 0.987788 |
| AKBR-6 | vWA | PfTRAP | 2.17E-08 | 1.36E+05 | 4.54E+03 | 2.96E-03 | 4.43E-05 | 0.024215 | 0.989348 |
| AKBR-7 | vWA | PfTRAP | 1.51E-10 | 3.17E+05 | 8.10E+03 | 4.78E-05 | 1.03E-05 | 0.113777 | 0.991841 |
| AKBR-10 | vWA | PfTRAP | 8.69E-09 | 1.51E+05 | 3.78E+03 | 1.31E-03 | 2.76E-05 | 0.028112 | 0.992658 |
| TY01 | vWA | PyTRAP | 1.65E-09 | 3.01E+05 | 7.22E+03 | 4.98E-04 | 2.21E-05 | 0.022956 | 0.989651 |
| TY02 | vWA | PyTRAP | 8.30E-10 | 1.24E+05 | 4.99E+03 | 1.03E-04 | 1.84E-05 | 0.185304 | 0.980355 |
| TY03 | vWA | PyTRAP | 1.65E-08 | 5.69E+05 | 4.96E+04 | 9.40E-03 | 1.93E-04 | 0.054834 | 0.973725 |
| TY04[#] | repeat | PyTRAP | *n/d* | *n/d* | *n/d* | *n/d* | *n/d* | *n/d* | *n/d* |
| TY05 | vWA | PyTRAP | 1.56E-08 | 1.83E+05 | 6.07E+03 | 2.86E-03 | 4.28E-05 | 0.02662 | 0.989553 |
| TY06 | vWA | PyTRAP | 3.95E-09 | 2.41E+05 | 7.32E+03 | 9.52E-04 | 3.01E-05 | 0.027236 | 0.988299 |
| TY07 | vWA | PyTRAP | 2.65E-08 | 1.50E+05 | 7.16E+03 | 3.97E-03 | 7.45E-05 | 0.027546 | 0.982021 |
| TY10 | vWA | PyTRAP | 5.82E-10 | 1.54E+05 | 5.37E+03 | 8.95E-05 | 1.55E-05 | 0.164387 | 0.988223 |
| TY11 | vWA | PyTRAP | 4.75E-09 | 1.13E+05 | 2.88E+03 | 5.37E-04 | 2.72E-05 | 0.055748 | 0.99251 |
| TY12 | vWA | PyTRAP | 2.20E-08 | 1.50E+04 | 3.42E+02 | 3.31E-04 | 1.56E-05 | 0.112929 | 0.989322 |
| TY13[#] | repeat | PyTRAP | *n/d* | *n/d* | *n/d* | *n/d* | *n/d* | *n/d* | *n/d* |
| TY14 | TSR | PyTRAP | 2.03E-07 | 2.25E+04 | 8.61E+02 | 4.58E-03 | 9.39E-05 | 0.038923 | 0.987681 |
| TY15 | TSR | PyTRAP | 7.95E-08 | 5.43E+03 | 1.27E+02 | 4.32E-04 | 1.54E-05 | 0.019626 | 0.993914 |
| TY19 | repeat | PyTRAP | 3.07E-07 | 1.61E+05 | 5.60E+04 | 4.95E-02 | 5.25E-03 | 0.07816 | 0.869796 |
| TY20 | vWA | PyTRAP | 1.03E-08 | 1.42E+05 | 4.21E+03 | 1.46E-03 | 3.35E-05 | 0.053002 | 0.988519 |

[&]Epitope assignments shown are based on data in Suppl. Table 3.
[*]Values shown are from global data fitting using a 1:1 binding model.
[#] Binding properties for TY04 and TY13 resulted in poor fits using the 1:1 binding model likely due to the presence of multiple epitopes per molecule of TRAP ectodomain, and were excluded from further kinetics analysis, see also Suppl. Fig. 7

*Supplementary Table 3: Coarse epitope mapping for anti-TRAP mAbs.*

| | | | ELISA signal for a given construct | | | | | |
|---|---|---|---|---|---|---|---|---|
| mAb name | epitope [a] | target | Ectodomain | vWA+TSR | TSR | TSR+repeat | vWA | repeats |
| AKBR-1 | TSR | PfTRAP | +++ | +++ | +++ | +++ | – | – |
| AKBR-3 | vWA | PfTRAP | ++ | + | – | – | – | – |
| AKBR-4 | vWA | PfTRAP | +++ | +++ | – | – | ++ | – |
| AKBR-5 | TSR | PfTRAP | +++ | – | +++ | +++ | – | – |
| AKBR-6 | vWA | PfTRAP | +++ | ++ | – | – | ++ | – |
| AKBR-7 | vWA | PfTRAP | ++ | ++ | – | – | ++ | – |
| AKBR-10 | vWA | PfTRAP | +++ | ++ | – | – | ++ | – |
| TY01 | vWA | PyTRAP | +++ | +++ | – | – | n/a | n/a |
| TY02 | vWA | PyTRAP | +++ | +++ | – | – | n/a | n/a |
| TY03 | vWA | PyTRAP | +++ | +++ | – | + | n/a | n/a |
| TY04 | repeat | PyTRAP | +++ | + | – | +++ | n/a | n/a |
| TY05 | vWA | PyTRAP | +++ | +++ | – | – | n/a | n/a |
| TY06 | vWA | PyTRAP | +++ | +++ | – | – | n/a | n/a |
| TY07 | vWA | PyTRAP | +++ | +++ | – | – | n/a | n/a |
| TY10 | vWA | PyTRAP | +++ | +++ | – | – | n/a | n/a |
| TY11 | vWA | PyTRAP | +++ | +++ | – | – | n/a | n/a |
| TY12 | vWA | PyTRAP | ++ | +++ | – | – | n/a | n/a |
| TY13 | repeat | PyTRAP | +++ | – | – | +++ | n/a | n/a |
| TY14 | TSR | PyTRAP | ++ | +++ | +++ | +++ | n/a | n/a |
| TY15 | TSR | PyTRAP | ++ | +++ | +++ | ++ | n/a | n/a |
| TY19 | repeat | PyTRAP | +++ | ++ | – | +++ | n/a | n/a |
| TY20 | vWA | PyTRAP | +++ | +++ | – | – | n/a | n/a |

[a] interpretation based on the results shown in each row

–: represents background value (≤ 2-fold ELISA background)

+: represents ≥5-fold ELISA background value

++: represents ≥10-fold ELISA background value

+++: represents ≥40-fold ELISA background value

n/a: test construct not available

*Supplementary Table 4. Monoclonal antibody sequence analysis and germline annotation.*

| mAb name | epitope& | target | IGHV# | IGHJ# | IGKV# | IGKJ# | CDRH3 length (aa)# | CDRL3 length (aa)# |
|---|---|---|---|---|---|---|---|---|
| AKBR-1 | TSR | PfTRAP | IGHV1S29*02 | IGHJ3*01 | IGKV3-4*01 | IGKJ2*01 | 9 | 9 |
| AKBR-3 | vWA | PfTRAP | IGHV3-2*02 | IGHJ2*01 | IGKV4-59*01 | IGKJ5*01 | 15 | 9 |
| AKBR-4 | vWA | PfTRAP | IGHV1-69*01,IGHV1-69*02 | IGHJ4*01 | IGKV17-127*01 | IGKJ5*01 | 14 | 8 |
| AKBR-5 | TSR | PfTRAP | IGHV6-6*01 | IGHJ4*01 | IGKV1-110*01 | IGKJ4*01 | 11 | 9 |
| AKBR-6 | vWA | PfTRAP | IGHV8-12*01 | IGHJ1*01 | IGKV12-46*01 | IGKJ1*01 | 15 | 9 |
| AKBR-7 | vWA | PfTRAP | IGHV1-9*01 | IGHJ3*01 | IGKV14-111*01 | IGKJ5*01 | 11 | 9 |
| AKBR-10 | vWA | PfTRAP | IGHV9-2-1*01 | IGHJ4*01 | IGKV4-57*01 | IGKJ4*01 | 14 | 9 |
| TY01 | vWA | PyTRAP | IGHV1S56*01 | IGHJ3*01 | IGKV3-4*01 | IGKJ1*01 | 7 | 9 |
| TY02 | vWA | PyTRAP | IGHV1-9*01 | IGHJ2*01 | IGKV3-4*01 | IGKJ2*01 | 6 | 9 |
| TY03 | vWA | PyTRAP | IGHV1S135*01 | IGHJ2*01 | IGKV6-25*01 | IGKJ5*01 | 13 | 9 |
| TY04 | repeat | PyTRAP | IGHV6-6*02 | IGHJ1*01 | IGKV8-30*01 | IGKJ1*01 | 11 | 9 |
| TY05 | vWA | PyTRAP | IGHV1-9*01 | IGHJ3*01 | IGKV3-4*01 | IGKJ4*01 | 6 | 9 |
| TY06 | vWA | PyTRAP | IGHV1-9*01 | IGHJ3*01 | IGKV3-4*01 | IGKJ1*01 | 6 | 9 |
| TY07 | vWA | PyTRAP | IGHV5-6-2*01 | IGHJ4*01 | IGKV3-12*01 | IGKJ2*01 | 11 | 10 |
| TY10 | vWA | PyTRAP | IGHV1-9*01 | IGHJ3*01 | IGKV3-4*01 | IGKJ2*01 | 6 | 9 |
| TY11 | vWA | PyTRAP | IGHV1-9*01 | IGHJ3*01 | IGKV3-4*01 | IGKJ2*01 | 6 | 9 |
| TY12 | vWA | PyTRAP | IGHV1-9*01 | IGHJ2*01 | IGKV5-43*01 | IGKJ1*01 | 10 | 9 |
| TY13 | repeat | PyTRAP | IGHV14-3*02 | IGHJ2*01 | IGKV5-43*01 | IGKJ5*01 | 12 | 9 |
| TY14 | TSR | PyTRAP | IGHV1-18*01 | IGHJ4*01 | IGKV6-15*01 | IGKJ5*01 | 8 | 9 |
| TY15 | TSR | PyTRAP | IGHV2-2*02 | IGHJ3*01 | IGKV10-96*01 | IGKJ1*01 | 13 | 7 |
| TY19 | repeat | PyTRAP | IGHV5-9-1*01 | IGHJ2*01 | IGKV19-93*01 | IGKJ1*01 | 12 | 8 |
| TY20 | vWA | PyTRAP | IGHV1-9*01 | IGHJ3*01 | IGKV3-4*01 | IGKJ2*01 | 6 | 9 |

&Epitope assignments shown are based on data in Suppl. Table 3.

### Assignments made from top-matching alleles using NCBI IgBLAST.

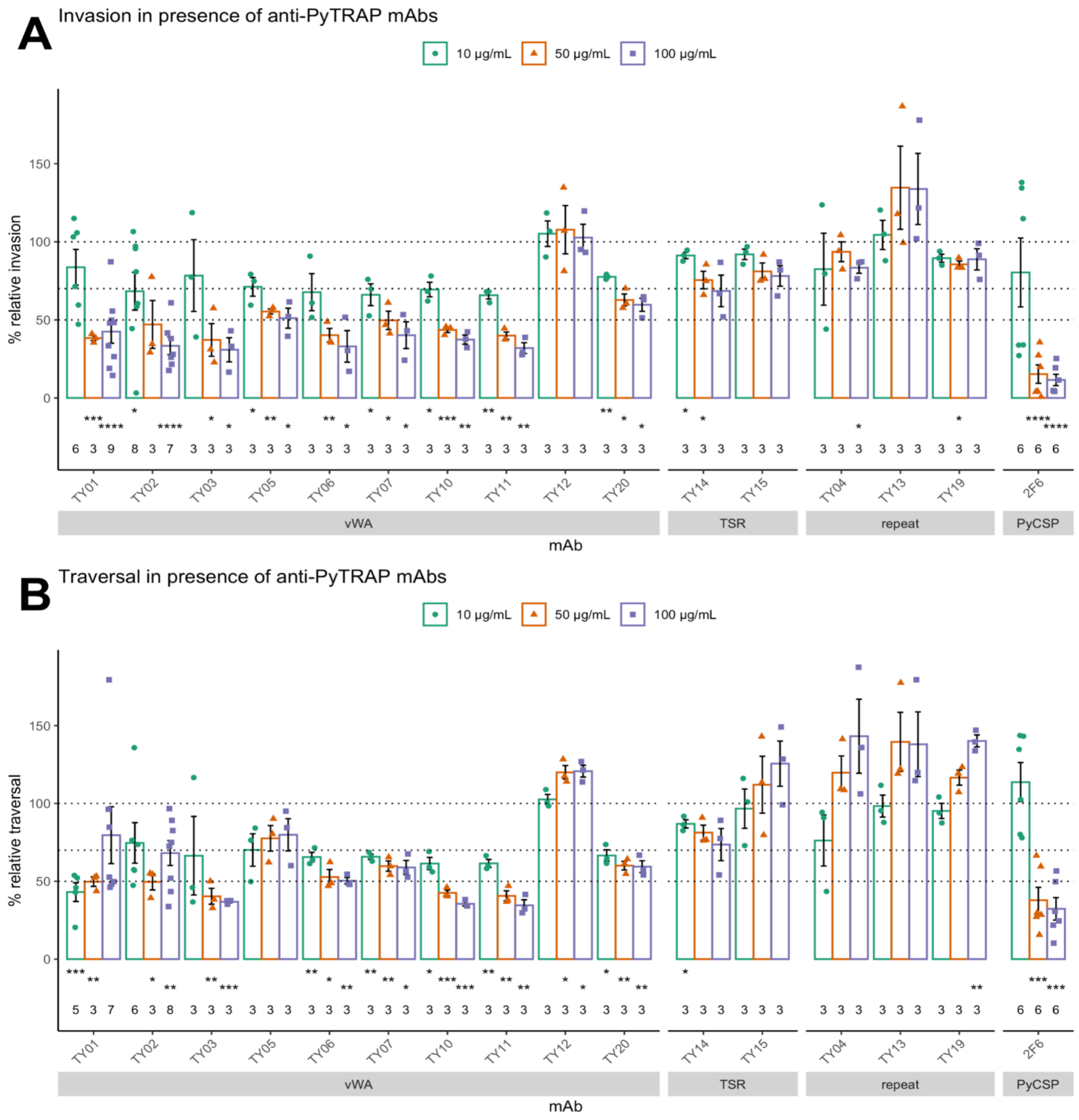


*Supplementary Figure 1. Monoclonal antibodies to PyTRAP inhibit parasite invasion, traversal in vitro.*

Each mAb was assessed for function in vitro for inhibition of invasion (**A**) and traversal (**B**). Each data point is an independent experiment showing the mean normalized % of non-specific mIgG-treated wells from technical duplicates. The number of independent replicate experiments is indicated below the corresponding bar. Asterisks indicate a significant difference from 100% as determined by two-tailed one-sample *t*-test: * is p≤0.05; ** is p≤0.01; *** is p≤0.001 and **** is p≤0.0001. Heat map showing the data for the 100-μg/mL condition is shown in **Fig. 3A**.

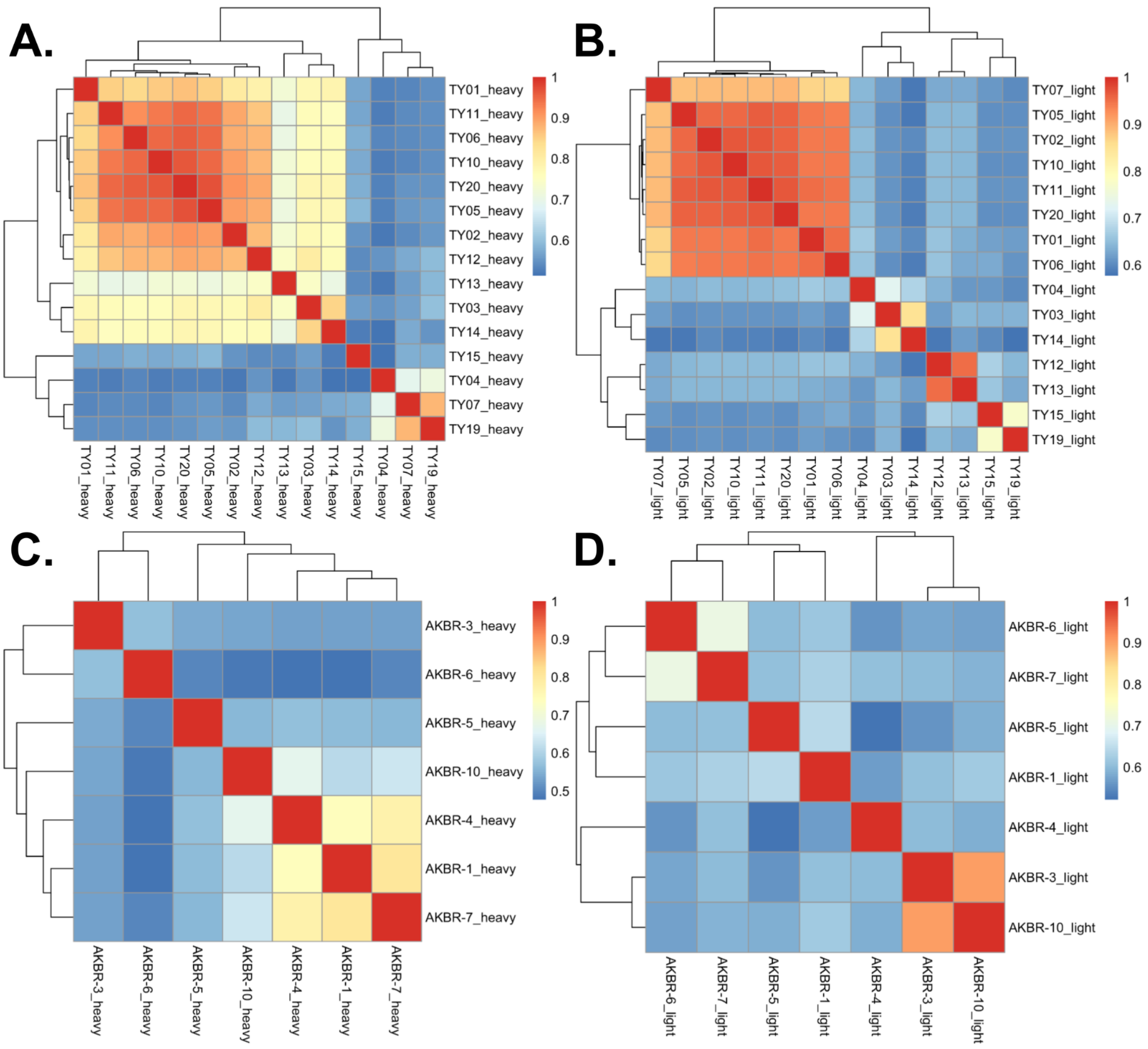


*Supplementary Figure 2. Sequence identity for heavy- and light-chain variable-segment sequences for anti-TRAP mAbs.*

Sequences for the variable-segments were aligned, each pairwise percent-identity values were determined and represented as a heat map color-coded according to the scale shown for each plot: (A) anti-PyTRAP heavy-chain sequences, (B) anti-PyTRAP light-chain sequences, (C) anti-PfTRAP heavy-chain sequences, (D) anti-PfTRAP light-chain sequences

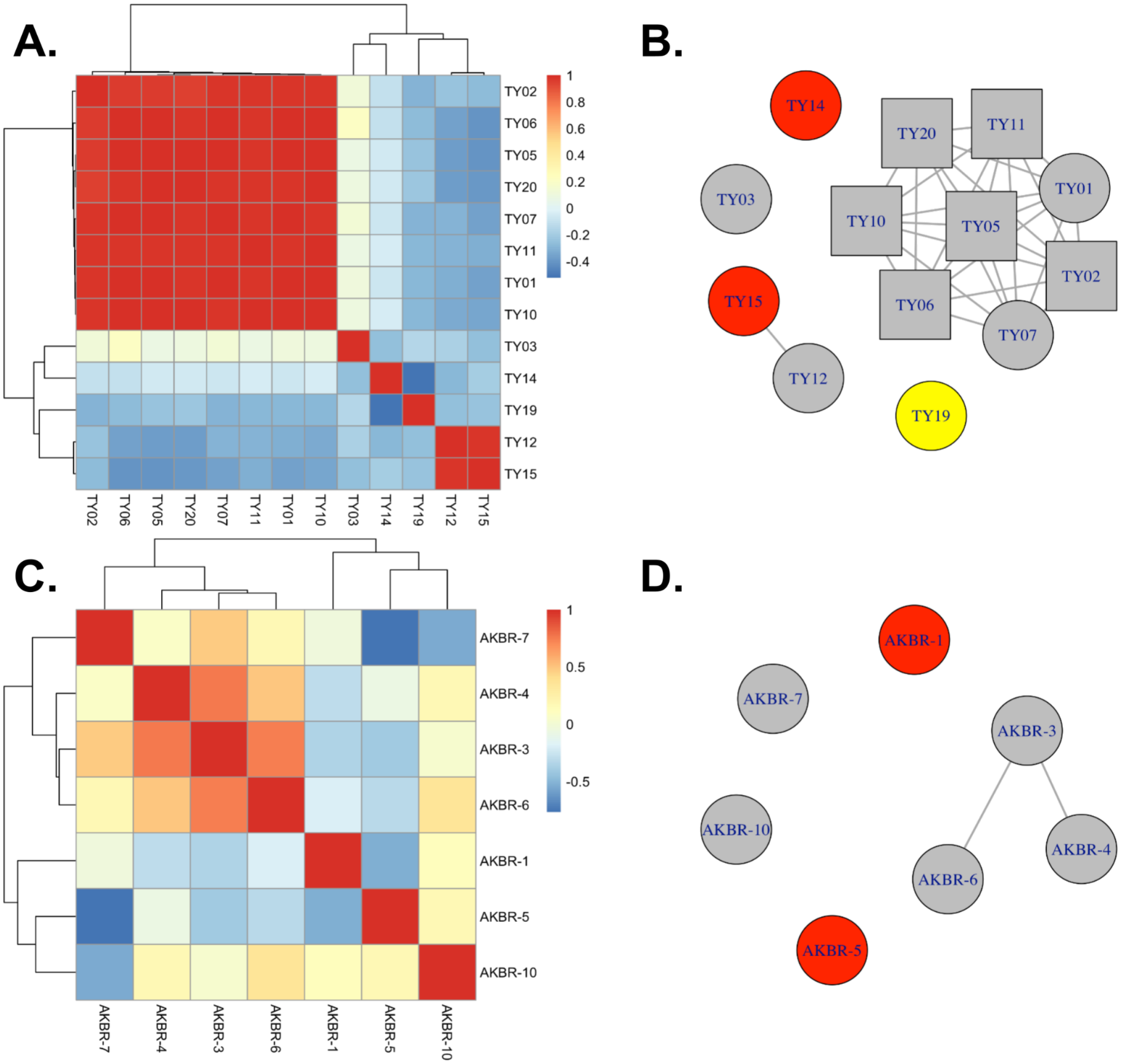


*Supplementary Figure 3. Epitope binning of anti-TRAP mAbs.*

(**A**) Clustered heatmap analysis representing Pearson correlation coefficients for interference patterns (i.e., degrees to which each mAb inhibited binding by the other mAbs in the panel) from pairs of anti-PyTRAP mAbs as measured by BLI.
(**B**) Network representation of the dataset shown in **A**. Edges connect pairs of nodes with a Pearson correlation coefficient >0.7. Nodes representing mAbs with high sequence identity (see Suppl. Fig. 2) are shown as squares with remaining nodes represented as circles. Gray fill is used for mAbs binding the vWA domain; red fill is used for mAbs binding the TSR domain; yellow fill is used for the mAb binding the repeat region. Clusters of interconnected nodes are referred to as bins.
(**C**) Clustered heatmap analysis representing Pearson correlation coefficients for interference profiles from pairs of anti-PfTRAP mAbs as measured by BLI.
(**D**) Network representation of the dataset shown in **C**. Edges connect pairs of nodes with a Pearson correlation coefficient >0.7. Gray fill is used for mAbs binding the vWA domain; red fill is used for mAbs binding the TSR domain. Clusters of interconnected nodes are referred to as bins.

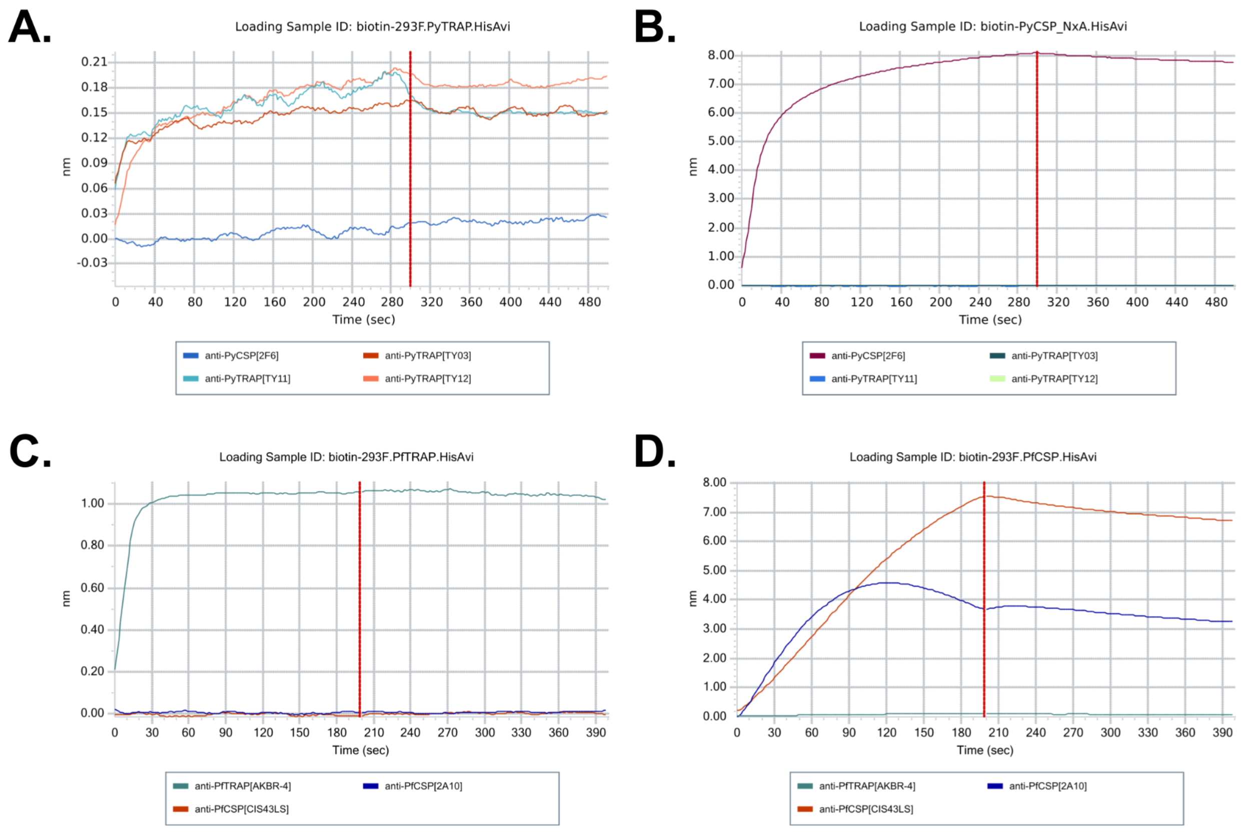


*Supplementary Figure 4. Assessment of cross-reactivity for mAbs used for passive immunization studies with purified recombinant antigens by BLI.*

(**A**) Anti-PyTRAP mAbs TY03, TY11 and TY12 were tested for binding to biotinylated recombinant PyTRAP alongside the anti-PyCSP mAb 2F6. (**B**) Anti-PyTRAP mAbs TY03, TY11 and TY12 were tested for binding to biotinylated recombinant PyCSP alongside the anti-PyCSP mAb 2F6. (**C**) Anti-PfTRAP mAb AKBR-4 was tested for binding to biotinylated recombinant PfTRAP alongside the anti-PfCSP mAbs 2A10 and CIS43LS (CIS43LS carries the M451L/N457S mutations in the Fc domain but has otherwise identical Ag-binding properties compared to the parental mAb CIS43). (**D**) Anti-PfTRAP mAb AKBR-4 was tested for binding to biotinylated recombinant PfCSP alongside the anti-PfCSP mAbs 2A10 and CIS43LS.

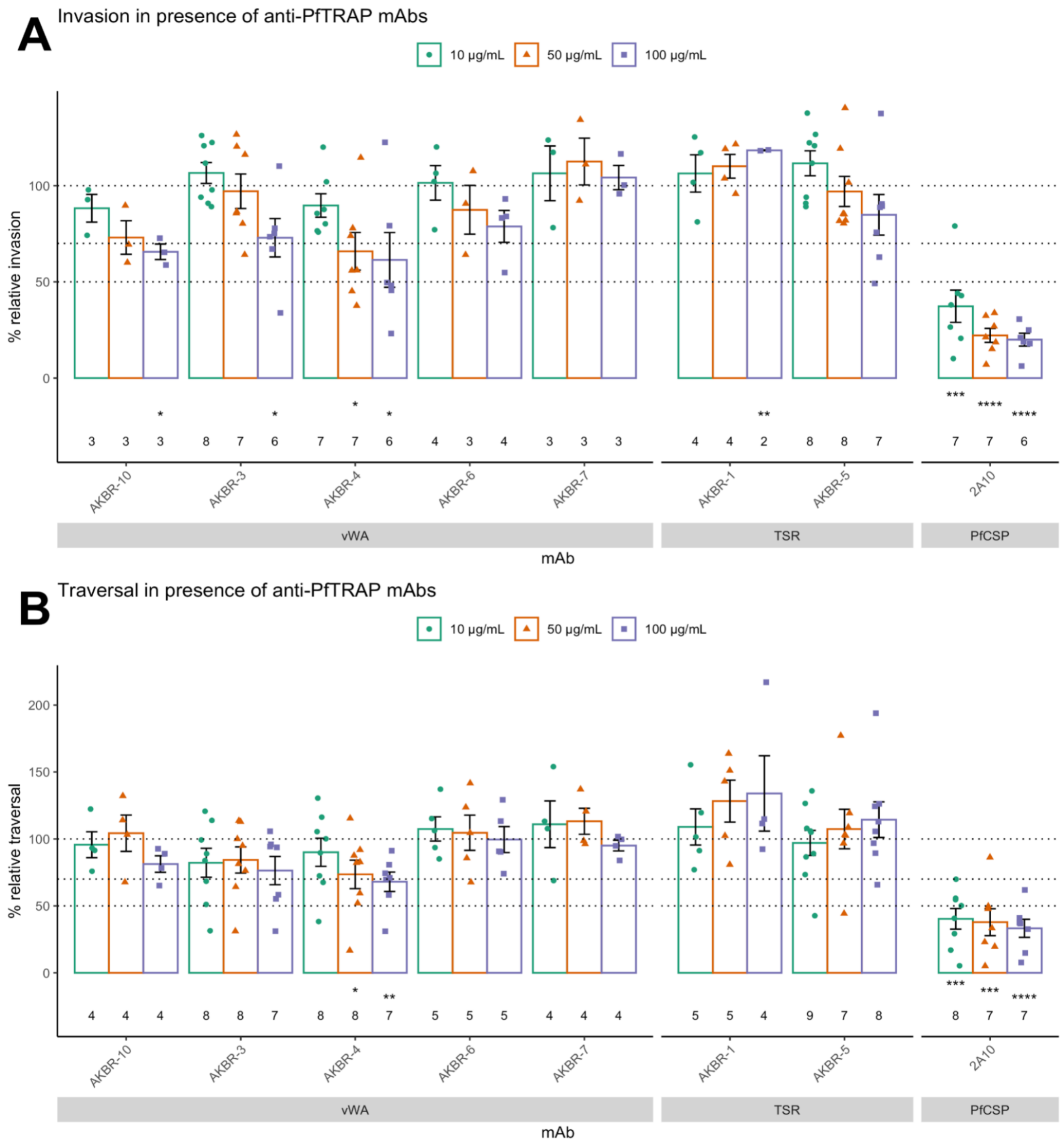


*Supplementary Figure 5. Monoclonal antibodies to PfTRAP inhibit parasite invasion and traversal in vitro.*

Each mAb was assessed for function in vitro for inhibition of invasion (**A**) and traversal (**B**). Each data point is an independent experiment showing the mean normalized % of non-specific mIgG-treated wells from technical duplicates. The number of independent replicate experiments is indicated below the corresponding bar. Asterisks indicate a significant difference from 100% as determined by two-tailed one-sample *t*-test: * is p≤0.05; ** is p≤0.01; *** is p≤0.001 and **** is p≤0.0001. Heat map showing the data for the 100-µg/mL condition is shown in **Fig. 5A**.

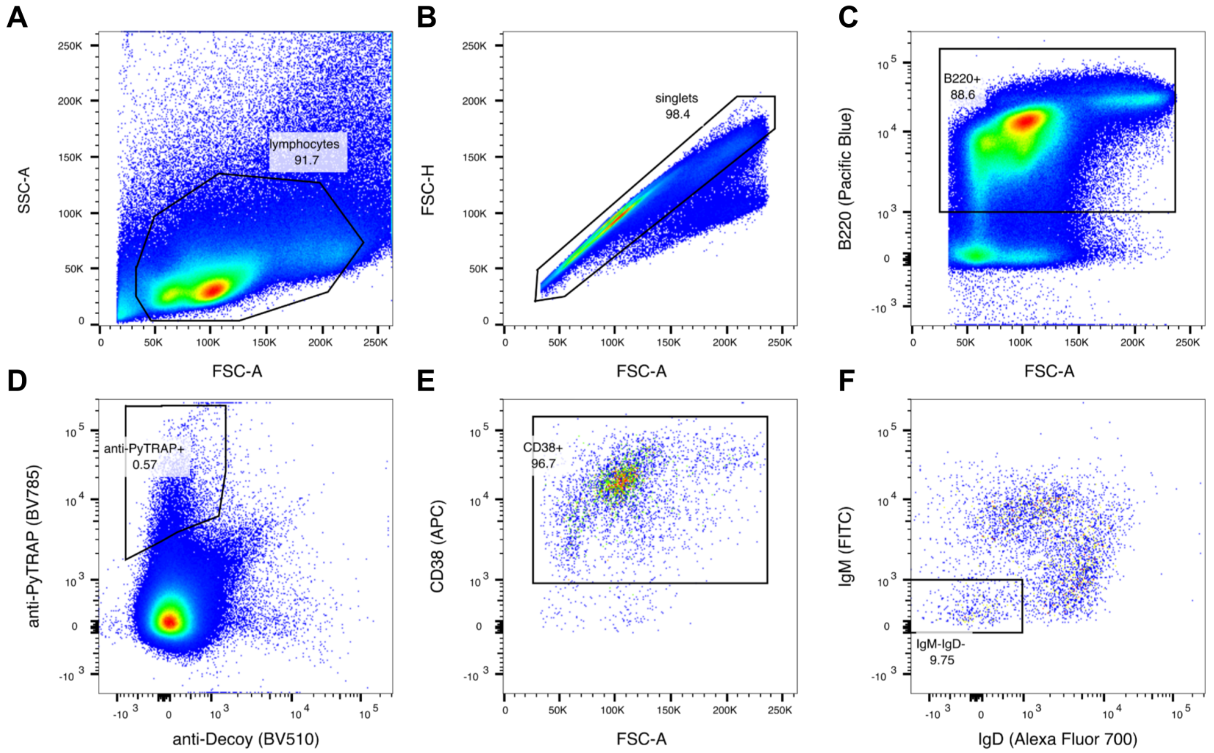


*Supplementary Figure 6. Sample gating strategy for isolating memory B cells for anti-TRAP mAb cloning.*

Splenocyte populations from immunized mice were identified and gated using "SSC vs FSC" plots (**A**). Single cells were then gated (**B**), followed by B220$^{+}$ populations (**C**), target-antigen-binding/decoy-antigen-free cells (**D**), CD38$^{+}$ cells (**E**) and IgM$^{-}$IgD$^{-}$ cells (**F**). Each pseudocolor plot representing marker stains has its axes labeled with the stain target and the fluorophore used, in parentheses.

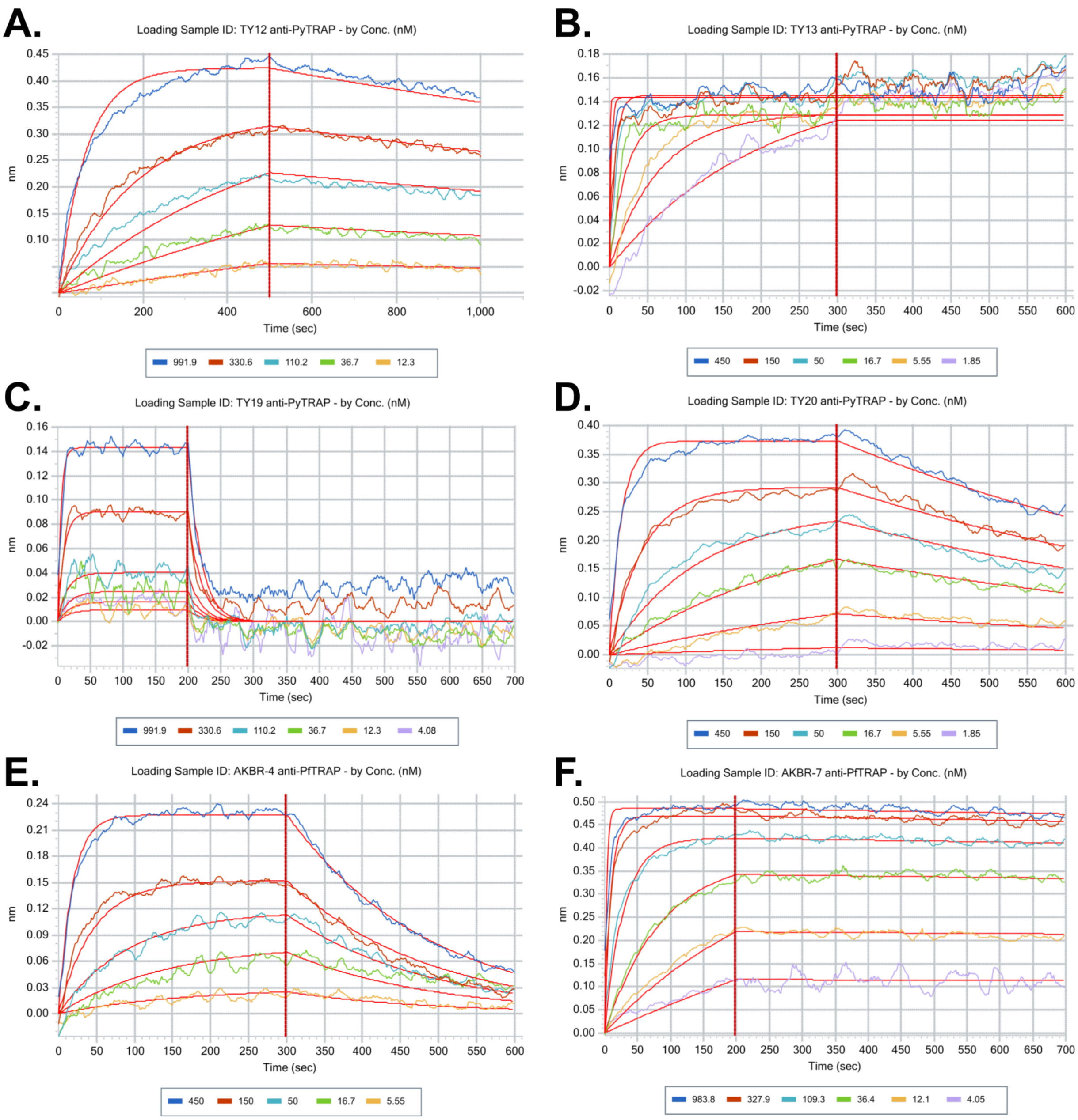


*Supplementary Figure 7. Sample data for measurements of mAb:antigen binding kinetics.*

Sensorgrams for multiple concentrations (color-coded according to the legends below each panel, nM) were used to generate a global fit for a 1:1 binding model (shown as red curves over the data). Transitions between association (the first 200–500 s) and dissociation (the last 300–500 s) phases are shown by a vertical red line. Representative data and their fits are shown: (**A**) TY12, (**B**) TY13 binding data did not result in a high-quality model fit likely due to recognition of multiple epitopes in the repeat region of PyTRAP, (**C**) TY19, (**D**) TY20, (**E**) AKBR-4, (**F**) AKBR-7.
